# Fungi hijack a plant apoplastic endoglucanase to release a ROS scavenging β-glucan decasaccharide to subvert immune responses

**DOI:** 10.1101/2021.05.10.443455

**Authors:** Balakumaran Chandrasekar, Alan Wanke, Stephan Wawra, Pia Saake, Lisa Mahdi, Nyasha Charura, Miriam Neidert, Milena Malisic, Meik Thiele, Murali Dama, Markus Pauly, Alga Zuccaro

**Author notes:** Shared first (Balakumaran Chandrasekar, Alan Wanke, Stephan Wawra).

## Abstract

Plant pathogenic and beneficial fungi have evolved several strategies to evade immunity and cope with host-derived hydrolytic enzymes and oxidative stress in the apoplast, the extracellular space of plant tissues. Fungal hyphae are surrounded by an inner, insoluble cell wall (CW) layer and an outer, soluble extracellular polysaccharide (EPS) matrix. Here we show by proteomics and glycomics that these two layers have distinct protein and carbohydrate signatures, implicating different biological functions. The barley (*Hordeum vulgare*) β-1,3-endoglucanase *Hv*BGLUII, which belongs to the widely distributed apoplastic glycoside hydrolase 17 family (GH17), releases a conserved β-1,3;1,6-glucan decasaccharide (β-GD) from the EPS matrices of fungi with different lifestyles and taxonomic positions. This low molecular weight β-GD does not activate plant immunity, is resilient to further enzymatic hydrolysis by β-1,3-endoglucanases due to the presence of three β-1,6-linked glucose branches and can scavenge reactive oxygen species. Additionally, exogenous application of β-GD leads to enhanced fungal colonization in barley. Our data highlights the hitherto undescribed capacity of this often overseen fungal EPS layer to act as an outer protective barrier important for fungal accommodation within the hostile environment at the apoplastic plant-microbe interface.

**Significance:** Here we identify and characterize a conserved β-1,3;1,6-glucan decasaccharide with antioxidant activity released from the fungal extracellular polysaccharide (EPS) matrix by the activity of a plant apoplastic endoglucanase. In addition, we provide a quantitative proteomic analysis of the fungal EPS and cell wall (CW) layers.

**HIGHLIGHTS:**

- The fungal extracellular polysaccharide (EPS) matrix and the cell wall (CW) are specific layers with distinct protein and carbohydrate signatures
- A conserved β-1,3;1,6-glucan decasaccharide (β-GD) is released from the EPS matrices of different fungi by the activity of the barley β-1,3-endoglucanase BGLUII, a member of the widely distributed apoplastic GH17 family
- The β-GD efficiently scavenges reactive oxygen species (ROS) and enhances fungal colonization
- The immunomodulatory potential as microbe-associated molecular pattern (MAMP) as well as the biochemical activity as ROS scavenger of soluble low molecular weight β-glucans are defined by the presence of β-1,6-glucose branches

## Introduction

The cell wall (CW) of animal pathogenic fungi is commonly surrounded by a soluble gel-like extracellular polysaccharide (EPS) matrix which often contains β-glucans, a heterogeneous group of glucose polymers (Gow *et al.*, 2017; Kang *et al.*, 2018). Commonly, fungal β-glucans have a structure comprising a main chain of β-1,3 and/or β-1,4-glucopyranosyl units, decorated by side chains with various branches and lengths (Han *et al.*, 2020). The immunomodulatory properties of β-glucans derived from animal pathogens have been long recognized (Goodridge *et al.*, 2009). β-glucans exhibit a broad spectrum of biological activities and have a dual role with respect to host immunity which depends on their chemo-physical characteristics. On the one hand, β-glucans are important microbe-associated molecular patterns (MAMPs) that are detected upon fungal colonization to trigger host immune responses in both vertebrates and invertebrates (Brown and Gordon, 2005). On the other hand, certain soluble β-glucans do not possess high immunogenic properties but are implicated in antioxidant activities and scavenging of reactive oxygen species (ROS) (Han *et al.*, 2020). Their immunological or antioxidant properties are rather complex and could be influenced by modifications in their structural characteristics such as molecular weight, substitution pattern, solubility, polymer charge and conformation in solution (Han *et al.*, 2020).

In plant-fungal interactions, carbohydrate metabolic processes mediated by carbohydrate-active enzymes (CAZymes) in the apoplast play a crucial role. Surface exposed and accessible fungal polysaccharides are hydrolyzed by apoplastic CAZymes, such as chitinases and glucanases and the resulting oligosaccharides can act as elicitors to trigger plant immune responses (Rovenich *et al.*, 2016). Recently, we described a soluble extracellular β-glucan matrix produced by endophytic fungi during root colonization of *Arabidopsis thaliana* (hereafter Arabidopsis) and barley (Wawra *et al.*, 2019). Very little is known about the biochemical properties, composition and function of this EPS matrix, but its detection in beneficial and pathogenic fungi strongly suggests a conserved role in counteracting environmental and immunological challenges during plant colonization (Wanke *et al.*, 2021). It is therefore crucial to investigate the structure and function of these soluble β-glucans and the hydrolytic events mediated by host apoplastic CAZymes during plant-fungal interactions. In this study, we have characterized the CW and the soluble EPS matrix produced by two distantly related fungi, the beneficial root endophyte *Serendipita indica* (Basidiomycota) and the pathogenic fungus *Bipolaris sorokiniana* (Ascomycota), spanning over 450 million years of evolution (Taylor and Berbee, 2006). Proteomics and glycomics revealed that β-glucan-binding proteins with WSC (cell <u>w</u>all integrity and <u>s</u>tress response <u>c</u>omponent) domains and β-1,3;1,6-glucan polysaccharides are enriched in the soluble EPS matrix compared to the CW layer. Treatment of the fungal EPS matrices with the apoplastic barley β-1,3-endoglucanase *Hv*BGLUII released a β-1,3;1,6-glucan decasaccharide (β-GD) with a mass/charge (m/z) of 1661 Da. Proton nuclear magnetic resonance (1H NMR) of β-GD is consistent with a heptameric β-1,3-glucan backbone substituted with three monomeric β-glucosyl residues at O-6. The β-GD is resilient to further enzymatic digestion by glycoside hydrolase 17 (GH17) family members and is immunologically inactive in a β-1,6-glucan side branch-dependent manner. This low molecular weight soluble β-GD is able to efficiently scavenge ROS and to enhance fungal colonization, confirming its role as a carbohydrate-class effector. The release of a conserved β-GD from the β-glucan-rich EPS matrix of a beneficial and a pathogenic fungus indicates that the utilization of this outermost CW layer as a protective shield against oxidative stress is a common strategy of root-colonizing fungi to withstand apoplastic defense responses during colonization.

## Materials and methods

### Commercial enzymes

The commercial β-glucanases used in this study are the *Trichoderma harzianum* lysing enzymes (TLE, Sigma L1412), the *Helix pomatia* β-1,3-glucanase (Sigma 67138) and the *H. vulgare* β-1,3-endoglucanase (GH17 family *Hv*BGLUII, E-LAMHV in 50% glycerol, Megazyme). TLE or *H. pomatia* β-1,3-glucanase were prepared in a stock concentration of 1.25 mg/ml in respective buffers: TLE (2 mM sodium acetate, pH 5.0), *H. pomatia* β-1,3-glucanase (2 mM MES pH 5.0). The GH17A family β-1,3-endoglucanase from *Formosa agariphila* (*Fa*GH17a) and the GH30 family β-1,6-exoglucanase from *Formosa* sp. nov strain Hel1_33_131 (*Fb*GH30) were obtained from Prof. Dr. Jan-Hendrik Hehemann (Center for Marine Environmental Sciences, University of Bremen, Germany) at a concentration of 5 μg/ml in water (Becker *et al.*, 2017).

### FITC488 labelling and confocal microscopy

FITC488 labelling of *Si*WSC3 and *Si*FGB1 and confocal laser scanning microscopy was done as described previously (Wawra *et al.*, 2019) using the KPL kit with the following modification for *Si*WSC3 labelling: 20 min incubation time at 20°C with half of the recommended concentration. The reaction was stopped by adding 1 M TRIS (pH 7.5) to a final concentration of 50 mM and then dialyzed overnight against 3 litres of Milli-Q water. Modifications to the standard labelling protocol were necessary because over labelling reduced the ability of the *Si*WSC3 to bind to its substrate (Wawra *et al.*, 2019).

### Microbial strains and culture conditions for EPS matrix production in *S. indica* and *B. sorokiniana*

The EPS matrix was isolated from the *S. indica* GFP strain in the dikaryotic wild-type background DSM11827 (Wawra *et al.*, 2016) and from the *B. sorokiniana* wild-type strain ND90r. *S. indica* spores were isolated from three-week-old cultures grown on solid complex medium (*Si*_CM) using 0.002% tween water (Zuccaro *et al.*, 2011). For the preculture, 2 ml of *S. indica* spores at a concentration of 500000/ml were inoculated in 100 ml of tryptic soy broth medium (TSB) containing 1% sucrose and shaking at 120 rpm at 28°C. After 48 hours the pre-cultures were transferred to 400 ml of TSB containing 1% sucrose and shaking at 120 rpm at 28°C for 72 hours. *B. sorokiniana* spores were isolated from 10 days old cultures grown on solid complete medium (*Bs*_CM), using 0.002% tween water (Sarkar *et al.*, 2019). The spores were inoculated at a final concentration of 250 spores/ml in 250 ml of yeast-extract peptone dextrose (YPD) medium shaking at 28°C for 36 hours.

### Isolation of EPS matrix from *S. indica* and *B. sorokiniana* culture supernatant

Culture supernatants from axenically grown *S. indica* or *B. sorokiniana* were collected by filtering the mycelia using Miracloth (Merck Millipore). The EPS matrix was isolated from the culture media using cryogelation. Briefly, the culture media were frozen overnight at −20°C and slowly thawed at room temperature for 16 hours. The precipitated EPS matrix (Figure S1) present in the culture medium was isolated using a pipette controller and washed four times with 30 ml of Milli-Q water and either used for proteome analyses (see next chapter) or washed one more time with 30 ml of 8.3 mM EDTA pH 8.0 to remove metal ions potentially present in the EPS. The proteins present in the EPS matrix were removed by treatment with 30 ml of protein denaturation solution (containing 8 M urea, 2 M thiourea, 4% SDS, 100 mM Tris-HCl pH 7.5) and boiling at 95°C for 15 minutes. The SDS present in the EPS matrix was removed by boiling the material with 30 ml of Milli-Q water at 95°C for 10 minutes and centrifuging at 11000 rpm for 10 minutes. The latter step was repeated approximately fifteen times until no further foaming occurred. The resulting protein-free EPS matrix was lyophilized and used for glycosyl linkage, thin-layer chromatography (TLC) or MALDI-TOF analyses.

### Proteome analysis of *S. indica* EPS matrix, CW and culture filtrate

The proteins were isolated from the EPS matrix, CW and the culture filtrate obtained from axenic cultures of *S. indica* GFP strain grown in different media (CM, YPD and TSB).

#### Protein isolation from the CW

Mycelium collected from the *S. indica* GFP strain was ground in liquid N_2_ and resuspended in PBS buffer containing 1 mM PMSF and 1% NP-40 using an ULTRA-TURRAX (IKA, Staufen, Germany). The resuspended mixture was incubated at 4°C in a rotating wheel for 30 minutes. The pellet obtained after centrifugation (8000xg) was resuspended in PBS buffer containing 1 mM PMSF and 0.1% IGEPAL using an ULTRA-TURRAX. The pellet obtained after centrifugation (8000xg) was washed three times with Milli-Q water. Finally, the pellet was resuspended in Laemmli buffer containing 8 M urea, 2 M thiourea and β-mercaptoethanol and boiled at 95°C for 10 minutes.

#### Protein isolation from the EPS matrix

The EPS matrix obtained from the culture media by cryogelation was washed four times with Milli-Q water and was directly boiled in Laemmli buffer containing 8 M urea, 2 M thiourea and β-mercaptoethanol at 95°C for 10 minutes.

#### Protein isolation from the culture filtrate

The culture supernatant left-over from the EPS matrix isolation step was first filtered using a Whatman filter paper and then using a 0.22 μM syringe filter. Approximately 30 ml of EPS matrix depleted culture supernatant were treated with 5 ml of 95% trichloroacetic acid and incubated overnight at 4°C. The precipitated proteins were collected by centrifugation (10000xg) for one hour at 4°C. The isolated proteins were washed at least three times with 100% acetone. The dried protein pellet was resuspended in Laemmli buffer containing 8 M urea, 2 M thiourea and β-mercaptoethanol and boiled at 95°C for 10 minutes.

### Liquid chromatography coupled mass spectrometric protein identification and quantification

The proteins isolated from the EPS matrix, CW and culture filtrate were separated using a 10% SDS PAGE gel for 15 minutes and subsequently stained with Coomassie Brilliant Blue. Protein-containing bands from Coomassie-stained gels were prepared for mass spectrometric analysis as described elsewhere (Poschmann *et al.*, 2014). Briefly, bands were destained and the proteins were reduced with dithiothreitol and alkylated with iodoacetamide and subjected to tryptic digestion. The resulting peptides were extracted and reconstituted in 0.1% (v/v) trifluoroacetic acid in water. Peptides were separated on an Ultimate 3000 Rapid Separation Liquid Chromatography system (RSLCS, Thermo Fisher Scientific) on a 25 cm length C18 column using a one-hour gradient and subsequently analysed by a QExactive Plus mass spectrometer (Thermo Fisher Scientific) as described with minor modifications (Poschmann *et al.*, 2014). First, survey scans were carried out at a resolution of 140000 and up to twenty-two- and threefold charged precursors selected by the quadrupole (4 m/z isolation window), fragmented by higher-energy collisional dissociation and fragment spectra recorded at a resolution of 17.500. Mass spectrometric data were further processed by MaxQuant 1.6.12.0 (Max-Planck Institute for Biochemistry, Planegg, Germany) with standard parameters if not otherwise stated. Label-free quantification, ‘match between runs’ and iBAQ quantification was enabled. Searches were carried out based on *S. indica* reference protein entries (UP000007148), downloaded on 15th Mai 2020 from the UniProt Knowledge Base. Carbamidomethylation at cysteines was considered as fixed and methionine oxidation and proteins N-terminal acetylation as variable modifications. Peptides and proteins were accepted at a false discovery rate of 1% and only proteins further considered were identified with at least two different peptides. The identified proteins were grouped into families using the Pfam database (Mistry *et al.*, 2021). The mass spectrometry proteomics data have been deposited to the ProteomeXchange Consortium *via* the PRIDE partner repository with the dataset identifier PXD025640.

### Large scale digestion and enrichment of the β-GD released from the EPS matrix of *S. indica* and *B. sorokiniana*

5 mg of freeze-dried EPS matrix obtained from *S. indica* or *B. sorokiniana* were ground with two stainless steel beads (5 mm) using a TissueLyser (Qiagen, Hilden, Germany) at 30 Hz for 1 minute at room temperature and soaked in 1 ml of 100 mM sodium acetate buffer pH 5.0 at 70°C overnight. The soaked material was incubated with 20 μl (50 units) of *Hv*BGLUII in a total reaction volume of 1 ml using 100 mM sodium acetate buffer pH 5.0. The digestion was performed at 40°C with shaking at 500 rpm for 48 hours. The supernatant containing the β-GD was collected by centrifugation at 10000xg for 5 minutes. The digested pellet was additionally suspended in 1 ml of water and heated at 80°C for 10 minutes to solubilize additional β-GD. The resulting supernatant was combined with the initial digested material and boiled at 95°C for 15 minutes. The precipitated proteins were removed by centrifugation and using a 0.45 μM syringe filter. The clear supernatant fraction was freeze-dried. The freeze-dried material was dialyzed against 3 liters of Milli-Q water at 4°C using a 1 kDa cut-off dialysis tubing (Repligen Spectra/Por 6 Pre-Wetted Regenerated Cellulose, cat. no. 888-11461). The dialyzate was lyophilized before further use.

### Preparation of alcohol insoluble residue and protein-free CW from *S. indica* and *B. sorokiniana*

The mycelium collected from axenic cultures of *S. indica* or *B. sorokiniana* was washed twice with Milli-Q water and freeze-dried overnight. The freeze-dried material was powdered in liquid N_2_ using a pestle and mortar and stored at −20°C until use. 20 mg of the material was resuspended in 1 ml of 70% aqueous ethanol with a stainless steel bead (5 mm) using a TissueLyser at 30 Hz for 1 minute. After centrifugation, the pellet was washed with chloroform/methanol (1:1), then acetone and subsequently air-dried. Protein-free fungal wall material was prepared as mentioned previously (Wawra *et al.*, 2016).

### Glycosyl linkage analysis of EPS matrices and CW preparations

EPS matrix (2 mg) or CW preparation of *S. indica* or *B. sorokiniana* were ground with a stainless steel bead (5 mm) using a TissueLyser mill at 30 Hz for 1 minute. The powdered material was subjected to glycosyl linkage analysis as described (Liu *et al.*, 2015). Briefly, a methylation reaction was performed using NaOH/DMSO. The methylated compounds were hydrolyzed in 1 M trifluoroacetic acid, reduced using sodium borodeuteride (ACROS Organics, cat.no. 194950050) and per-o-acetylated. The resulting partially methylated alditol acetates were analyzed using an Agilent 5977A GC/MSD System equipped with a SP-2380 Fused Silica Capillary Column (Supelco). The glycosidic linkages were assigned based on retention time and mass spectrum fragmentation patterns compared to the CCRC spectral database (https://www.ccrc.uga.edu/specdb/ms/pmaa/pframe.html).

### Digestion of EPS matrix and CW and TLC analysis

Freeze-dried EPS matrix (1 mg) or CW preparation (1 mg) from *S. indica* or *B. sorokiniana* were suspended in 400 μl of 2 mM sodium acetate pH 5.0 (for TLE), 2 mM MES pH 5.0 (for *H. pomatia* β-1,3-glucanase) or 100 mM sodium acetate (for *Hv*BGLUII) pH 5.0 at 70°C overnight. The excess buffer was removed and the suspended material was treated with 5 μl of the glucanase enzymes (0.125 mg/ml or 12.5 units for *Hv*BGLUII) in the respective buffers, containing 1 μl of BSA (100 mg/ml) in a total reaction volume of 50 μl and incubated at 40°C by shaking at 500 rpm for 16 hours. The digestion reaction was stopped by incubating the samples at 95°C for 10 minutes. An aliquot was subjected to TLC using a silica gel 60 F254 aluminum TLC plate (Merck Millipore, cat.no.105554,) with a running buffer containing ethyl acetate/acetic acid/methanol/formic acid/water in a ratio of 80:40:10:10:10 (v/v). D-glucose, laminaribiose β-1-3-(Glc)_2_, laminaritriose β-1-3-(Glc)_3_, gentiobiose β-1-6-(Glc)_2_ and laminaripentaose β-1-3-(Glc)_5_ with a concentration of 1.5 mg/ml were used as standards. The glucan fragments were visualized by spraying the TLC plate with glucan developer solution (containing 45 mg N-naphthol, 4.8 ml H_2_SO_4_, 37.2 ml ethanol and 3 ml water) and heating the TLC plate to 100°C until the glucan bands are visible (approximately 4-5 minutes).

### MALDI-TOF analysis

#### Analysis of the β-GD from S. indica

Freeze-dried β-GD was solubilized in water at 70°C for 10 minutes.

#### Structural characterization of the β-GD from S. indica and B. sorokiniana

10 μl of β-GD (2 mg/ml) were treated with 2 μl of *Fa*GH17a or 2 μl *Fb*GH30 or 1 μl of *Fa*GH17a + 1 μl *Fb*GH30 in 50 μl of Milli-Q water. The digestion reaction was carried out at 40°C for 16 hours and the reaction was stopped by incubation at 95°C for 5 minutes.

#### Analysis of oligosaccharides released from the EPS matrix and CW of S. indica

Freeze-dried EPS matrix (1 mg) or CW preparation (1 mg) isolated from *S. indica* were suspended in 400 μl of a 2 mM sodium acetate buffer, pH 5.0 (TLE), 2 mM MES buffer, pH 5.0 (*H. pomatia* β-1,3-glucanase), or 25 mM sodium acetate buffer, pH 5.0 (*Hv*BGLUII) and incubated at 70°C overnight. The suspended material was treated with 2.5 μl of TLE, 2.5 μl of *H. pomatia* β-1,3-glucanase or 1 μl of *Hv*BGLUII in the respective buffers, as described before, in a total reaction volume of 50 μl. The digestion was performed at 40°C with shaking at 500 rpm for 16 hours. The digestion reaction was stopped at 95°C for 10 minutes and centrifuged at 11000xg.

#### Analysis of oligosaccharides released from the EPS matrix and CW of B. sorokiniana

Freeze-dried EPS matrix (1 mg) or CW preparation (1 mg) isolated from *B. sorokiniana* were solubilized in 400 μl of 2 mM sodium acetate, pH 5.0 (TLE) or 25 mM sodium acetate, pH 5.0 (*Hv*BGLUII). The solubilized material was treated with 1 μl of TLE or 1 μl of *Hv*BGLUII in a total reaction volume of 50 μl. The digestion was performed at 40°C with shaking at 500 rpm for 16 hours. The digestion reaction was stopped at 95°C for 10 minutes.

#### Mass spectrometrical analysis

The oligosaccharides present in the prepared samples were analysed by Oligosaccharide Mass Profiling as described (Günl *et al.*, 2011). Briefly, the samples were spotted onto a dried spot of dihydroxy benzoic acid matrix (10 mg/ml) and analysed by matrix-assisted laser desorption ionization – time of flight mass spectrometry (Bruker rapifleX instrument). The machine was set to linear, positive reflectron mode with an accelerating voltage of 20000 V. The spectra from the samples were analysed using flexanalysis software 4.0 (Bruker Daltonics).

### Reduction and purification of *S. indica* β-GD

Enriched β-GD (40 mg) was reduced with sodium borodeuteride (20 mg/ml) (ACROS Organics, cat.no. 194950050) in 1 M ammonium hydroxide for 90 minutes at room temperature. The reaction was neutralized by addition of glacial acetic acid and 9:1 (v:v) methanol:acetic acid. The solvents were evaporated under N_2_ gas. The dried material was washed once with 9:1 (v:v) methanol:acetic acid and three times with methanol. In each washing step, the methanol:acetic acid or methanol were evaporated under N_2_ gas. The dried material was dissolved in 6% aqueous methanol, vortexed and centrifuged to remove any occurring debris. The supernatant (50 μl) was subjected to reverse-phase chromatography using a Vydac 238 TP C18 column (Vydac, Hesperia, CA, USA) eluting with a linear gradient from 6% to 12% methanol in 10 minutes, followed by 12% to 50% methanol in 10 minutes and equilibrated back to 6% methanol in 10 minutes with a flow rate of 0.5 ml/min. The eluting compounds were detected by an evaporative light scattering detector (ERC GmbH, Munich, Germany) at 38°C and simultaneously collected for MALDI-TOF analyses. Collected fractions containing the β-GD were pooled and freeze-dried.

### ^1^H NMR analysis

Reduced β-GD (1 mg) was dissolved in D_2_O (100% atom D, ACROS Organics, 320700075) at 80°C for 10 minutes and subsequently freeze-dried overnight. The freeze-dried material was dissolved in 300 μl of 6:1 (v/v) of methyl sulphoxide D6 (99.9% atom D + 1% tetramethylsilane, ACROS Organics):D_2_O (100% atom D, ACROS Organics, 320700075) at 80°C for 10 minutes. The ^1^H NMR spectrum of the reduced β-GD was measured using a 600 MHz Bruker NMR spectrometer at 80°C (Kim *et al.*, 2000). The chemical shift signal was referred to the internal standard tetramethylsilane at 0 ppm and the ^1^H NMR spectrum was processed using Bruker’s Topspin software.

### Purification of native β-GD for the ROS burst assay

Enriched β-GD (40 mg) was dissolved in 6% aqueous methanol, vortexed and centrifuged to remove any debris. The β-GD was purified as described above but without reduction and used for the ROS burst assays if not otherwise stated in the legend.

### Plant material and growth conditions

Immunity assays were performed with barley (*H. vulgare* L. cv Golden Promise) and *thaliana* Col-0. Barley seeds were surface sterilized with 6% bleach for one hour, followed by five washing steps with 30 ml of sterile Milli-Q water (each 30 minutes, two quick rinses for a couple of seconds between longer incubation steps). Sterilized seeds were germinated for three days on wet filter paper at room temperature in the dark under sterile conditions. Seedlings were transferred to sterile jars containing solid 1/10 PNM (plant nutrition medium, 0.4% gelrite [Duchefa, Haarlem, the Netherlands]) (Lahrmann *et al.*, 2013) and cultivated in a growth chamber with a day/night cycle of 16/8◻h (light intensity, 108◻μmol◻ m^−2^ sec^−1^) and temperatures of 22/18°C. Seedlings were grown for four more days before being used for immunity assays.

Arabidopsis seeds were surface sterilized (10 min 70% ethanol, 7 min 100% ethanol) and sown on ½ Murashige & Skoog (MS) medium (pH 5.7) supplemented with 0.5% sucrose and 0.4% Gelrite (Duchefa, Haarlem, the Netherlands). Plates were transferred to climate chambers with 8/16 h light/dark regime (light intensity of 110 μmol m^−2^ sec^−1^) at 22/18°C. Seven day old seedlings were transferred into petri dishes filled with 30 ml fresh ½ MS liquid medium and grown for seven additional days under the same conditions.

### Preparation of crude fungal CW substrates for immunity assays

For crude extraction of soluble fragments from the fungal CW and EPS matrix, 5 mg of the respective fungal substrate were transferred to a 2 ml Eppendorf tube with three stainless steel beads (5 mm), snap-frozen in liquid nitrogen and ground with a TissueLyser (1 min, 30 Hz). The ground substrate was resuspended in 2 ml of Milli-Q water and incubated at 70°C for 16 hours (700 rpm), then boiled at 95°C for 10 minutes. The volume was filled up to 5 ml for a final concentration of 1 mg/ml. If indicated, the substrate was additionally sonicated for 2 min (output control 5, duty cycle 30%) using a Branson Sonifier 250 (Branson Ultrasonics, Brookfield, CT, USA). The suspension was centrifuged at 13000xg for 10 minutes and the supernatant was further used in plant immunity assays.

### Plant immunity assays

For immunity assays using barley, roots were separated from seven-day-old seedlings (cut 2 cm below seed), root tips were removed (first 1 cm) and residual roots were cut into 5 mm pieces. Each assay was performed with randomized root pieces from 16 barley seedlings. Four root pieces were transferred into each well of a 96-well plate microtiter plates containing Milli-Q water. For immunity assays with Arabidopsis, intact 14-day-old seedlings were transferred into a 96-well plate (1 seedling/well) filled with Milli-Q water. Plant material was incubated overnight at room temperature on the bench. The next day, water was replaced by 100 μl of fresh Milli-Q water containing 20 μg/ml horseradish peroxidase (Sigma-Aldrich, Taufkirchen, Germany) and 20 μM L-012 (Wako Chemicals, Neuss, Germany). Following a short incubation time (approx. 15 min), 100 μl of double-concentrated elicitor solutions were added to the wells. Substrates from fungal CWs were prepared as described earlier. Chitohexaose was purchased from Megazyme (Bray, Ireland), flg22 from GenScript (Piscataway, NJ, USA) and laminarin from Sigma-Aldrich (Taufkirchen, Germany). All commercial substrates were dissolved in Milli-Q water without additional treatment. Measurements were started immediately and taken continuously with an integration time of 450 ms using a TECAN SPARK 10M. After the ROS burst assay (1 hour after elicitor addition), roots from 3-4 wells were pooled, dried on tissue paper and snap-frozen in liquid nitrogen for further analysis of gene expression changes. Extraction of total RNA was performed using TRIzol reagent (Invitrogen, Karlsruhe, Germany) and contaminating gDNA was digested during a DNaseI treatment (Thermo Fisher Scientific, Schwerte, Germany) according to manufacturers’ instructions. Synthesis of cDNA was carried out using the First Strand cDNA Synthesis Kit (Thermo Fisher Scientific, Schwerte, Germany) without changes to the manufacturer’s protocol. Target gene expression was analysed by quantitative reverse transcription PCR (RT-qPCR) as described previously (Wawra *et al.*, 2019). Relative expression of elicitor-responsive *HvWRKY2* gene (Shen *et al.*, 2007) compared to *H. vulgare* ubiquitin gene (Sarkar *et al.*, 2019) was determined using the following primer pairs: *HvUBI*_F (5′-ACCCTCGCCGACTACAACAT-3′) with *HvUBI*_R (5′-CAGTAGTGGCGGTCGAAGT-3′) and *HvWRKY2*_F (5′-AACAACCACCACCAGTCGTT-3′) with *HvWRKY2*_R (5′-TCACCTTCTGCCCGTACTTC-3′). Gene expression levels in elicitor-treated samples were normalized to the expression levels in mock-treated samples.

### In vitro 3,3’-diaminobenzidine (DAB) oxidation assay

DAB assay was performed as previously described (Nostadt *et al.*, 2020). In short, sugar substrates (concentrations as indicated) were mixed with 0.05 μM horseradish peroxidase (Sigma-Aldrich, Taufkirchen, Germany) and 1 mM H_2_O_2_ (Sigma-Aldrich, Taufkirchen, Germany) in reaction buffer (50 mM sodium acetate, 150 mM NaCl, pH 5) and incubated for 10 minutes at room temperature in a 96-well plate. Then, an equal volume of a 200 μM DAB solution was added to the wells. After 16 hours of incubation in the dark, plates were scanned on a flatbed scanner (transmissive light mode).

### Fenton reaction-based oxidation of sugars

The assay was performed as previously described (Matros *et al.*, 2015). In short, carbohydrate samples (300 μM) were mixed with Fenton reagents [1 mM H_2_O_2_ (Sigma), 100 μM FeSO_4_ (Sigma)] and incubated overnight at 30°C. Control samples with an additional 100 μM EDTA (Sigma) or without Fenton reagents were treated in the same way. Samples were centrifuged at 13000xg for 10 min and the supernatant was further analysed by MALDI-TOF.

### Colonization assay

Roots of four-day-old barley seedlings were inoculated with 3 ml of *S. indica* spores at a concentration of 500000/ml and grown at a day/night cycle of 16/8 h at 22/18°C, 60% humidity and 108 μmol m^−2^ sec^−1^ light intensity. At one and two days post inoculation (dpi), 1 ml of sterile water as a control or 100 or 300 μM β-GD were added to the jars which contained 4 seedlings each. The seedlings of each jar were pooled and harvested at 3 dpi. The roots were washed in ice water to remove extraradical hyphae, cut as previously described (Nizam *et al.*, 2019), frozen in liquid nitrogen and used for RNA extraction. RNA extraction for fungal colonization, cDNA generation and RT-qPCR were performed as previously described (Sarkar *et al.*, 2019). For quantification of fungal colonization by RT-qPCR, the following primers were used: 5’-GCAAGTTCTCCGAGCTCATC-3’ and 5’-CCAAGTGGTGGGTACTCGTT-3’ for *S. indica* translation-elongation factor (*SiTEF*) and 5’-ACCCTCGCCGACTACAACAT-3’ and 5’CAGTAGTGGCGGTCGAAGTG3’ for barley ubiquitin (*HvUbi*).

## Results

### Beneficial and pathogenic fungi produce a gel-like β-glucan EPS matrix surrounding their CW

We recently reported on a gel-like EPS matrix surrounding the hyphae of different fungi during colonization of plant hosts (Wanke *et al.*, 2021; Wawra *et al.*, 2019), suggesting that secretion of these soluble glycans is a common feature of root-associated fungi independent of their lifestyle and taxonomy. This finding motivated us to investigate the biochemical characteristics, composition and function of the matrix in the beneficial root endophyte *S. indica* and in the pathogenic fungus *sorokiniana*. To this end, we first labelled the β-1,3-glucan-binding lectin PIIN_05825 (*Si*WSC3-His-FITC488) from *S. indica* by applying an improved FITC488 conjugation protocol (see methods) and used it as a molecular probe for localization studies of the fungal EPS matrix *in planta*. Both fungi produced an EPS matrix containing β-1,3-glucans surrounding the WGA (wheat germ agglutinin)-stained chitin layer of fungal hyphae (Figure 1). Consistent with our previous observations, the localization of the specific β-1,6-glucan-binding lectin *Si*FGB1 (Wawra *et al.*, 2016) to this matrix indicates that β-1,6-linked glucans are also present (Figure 1). This expands the repertoire of fungal β-glucan-binding lectins that can be used as molecular probes to fluorescently label the fungal EPS matrix during live cell imaging.

**Figure 1.**
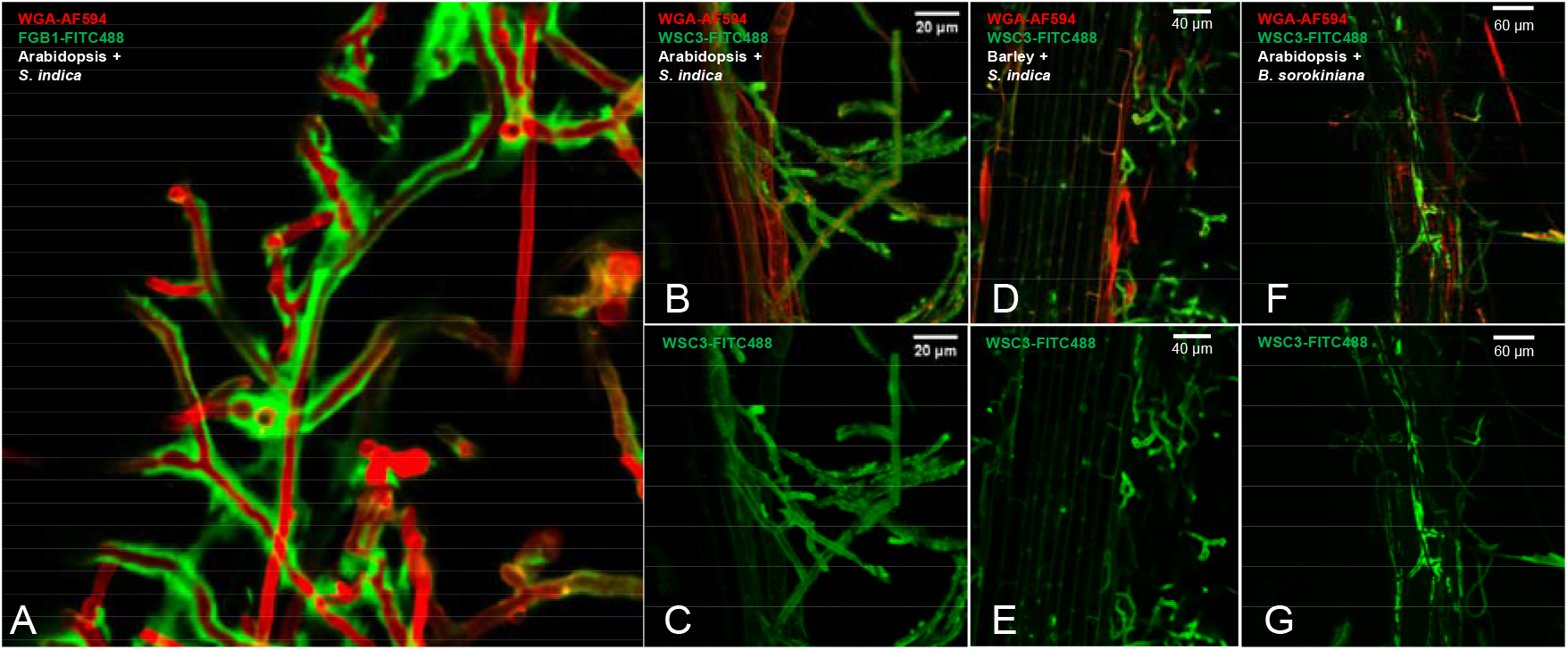
Fungal EPS matrix revealed by fluorescently labelled β-glucan binding lectins and confocal laser scanning microscopy during root colonization. The β-glucan-binding *Si*FGB1 or *Si*WSC3-His and the chitin-binding WGA lectins were used as molecular probes to visualize the fungal EPS matrix and CW of *S. indica* and *B. sorokiniana*. Green fluorescence corresponds to FITC488-labelled *Si*FGB1 or *Si*WSC3-His. Red fluorescence corresponds to WGA-AF594. Panels A, B, D and F are merged confocal microscopy images of *Si*FGB1-FITC488 or *Si*WSC3-His-FITC488 and WGA-AF594. Panels C, E and G display the EPS layer of *S. indica* or *B. sorokiniana* stained by *Si*WSC3-His-FITC488 during colonization of Arabidopsis or barley roots. Bars, 20 μm (panels: B, C), 40 μm (panels: D, E), 60 μm (panels: F, G). CW: cell wall; EPS: extracellular polysaccharide; WSC: cell wall integrity and stress response component; WGA: wheat germ agglutinin.

### The *S. indica* EPS matrix, CW and culture filtrate represent three functionally distinct but interconnected compartments

To identify secreted proteins associated with the EPS matrix, the CW and/or the culture filtrate, we performed comparative proteomics with protein extracts from these three compartments from *S. indica* axenically grown in three different media (CM, YPD, TSB). We identified 1724 proteins from all media and compartments (Table S1, Figure 2A). Among those, 220 proteins carried a predicted signal peptide, which were further analysed for their domain architecture using the Pfam database (Table S2 and Figure 2A). Glucan-binding proteins with at least one WSC domain were highly enriched or uniquely present in the EPS matrix compared to the CW or the culture filtrate, irrespective of the medium used (Figure 2B and Figure S2). Four of them, including *Si*WSC3, represented the most abundant proteins in this compartment (Figure 2B and Figure S2), thereby confirming our *in planta* localization study (Figure 1B-G). Gene expression analyses revealed an induction of these genes during *S. indica* colonization of barley and Arabidopsis over time (Wawra *et al.*, 2019) (Figure S3). Additionally, several proteases and CAZymes were enriched in the EPS matrix and culture filtrate but not in the CW (Figure S2), suggesting that the matrix may serve as a transient storage depot for these proteins. In the CW fraction, we identified two chitin-binding LysM proteins (PIIN_02172 and PIIN_02169) and several other lectin-like proteins, including the β-1,6-glucan-binding effector *Si*FGB1 (Wawra *et al.*, 2016) (PIIN_03211) and the ricin B lectin (PIIN_01237). Most of these lectins were also present in the culture filtrate, indicating that besides their ability to bind to CW components, such as chitin and glucan, they also function as soluble lectins in the extracellular environment (Figure 2B and Figure S2). This is in agreement with the function of *Si*FGB1, which has the potential to alter fungal CW composition and properties as well as suppress β-glucan-triggered immunity in the apoplast of different plant hosts (Wawra *et al.*, 2016). Remarkably, proteins with the cellulose-binding CBM_1 or the starch-binding CBM_20 domains were enriched in the culture filtrate (Figure S2). Based on the distribution and nature of the carbohydrate-binding proteins we conclude that the EPS matrix, the CW and the culture filtrate represent three functionally distinct but interconnected compartments.

**Figure 2.**
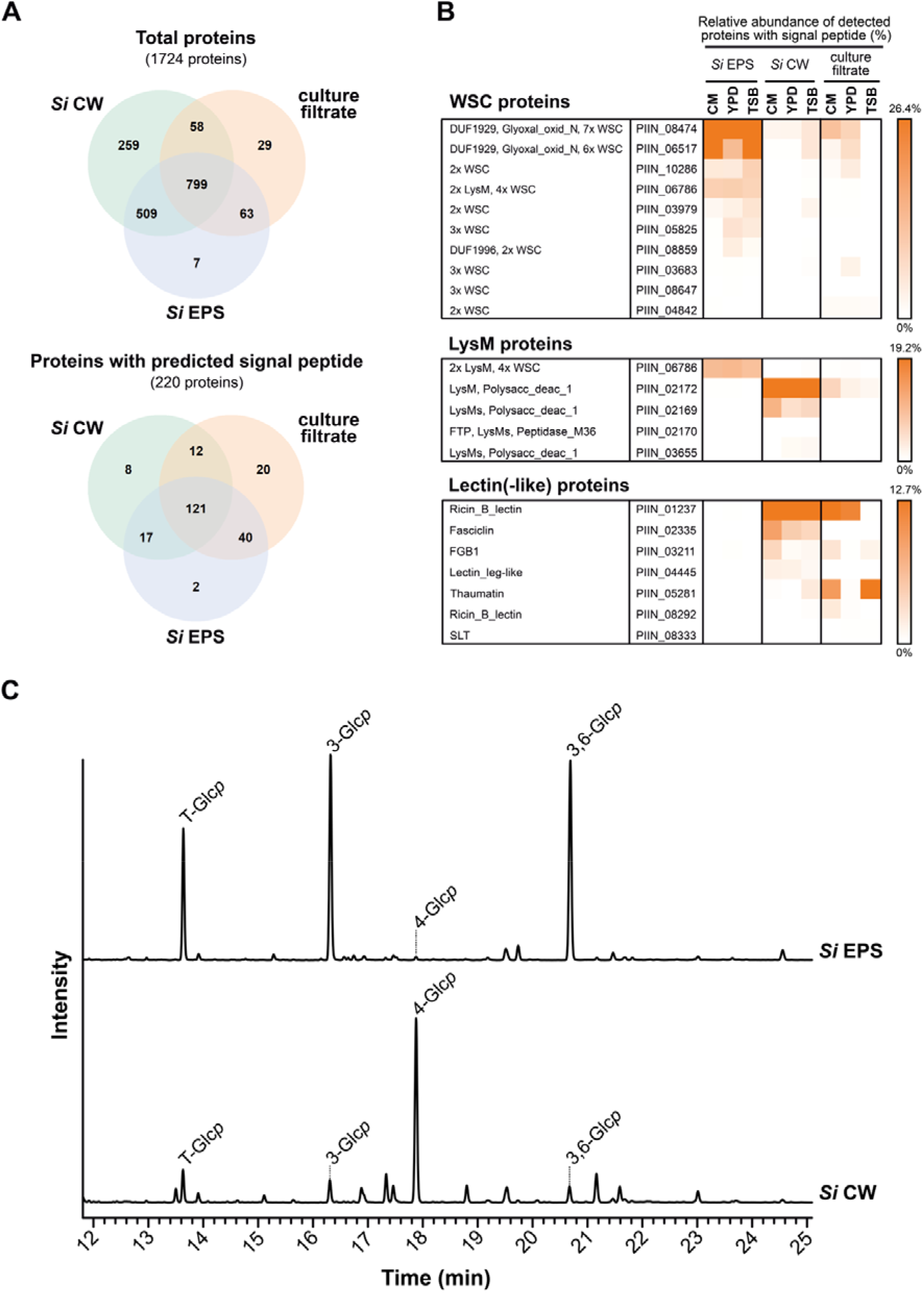
Proteomics and glycosyl linkage analysis of *S. indica* EPS matrix and CW. (A) Venn diagram of proteins identified in the EPS matrix, CW and/or culture filtrate from three cultivation media (CM, YPD and TSB). (B) Proteins with WSC domain/s are enriched in the EPS matrix of *S. indica* (see also Figure S2). The relative abundance of each protein was calculated using label-free quantification intensity values and is depicted in percentage. (C) Glycosidic linkage analysis of *S. indica* EPS and CW preparations. 3-glucose and 3,6-glucose are abundant in the EPS matrix, whereas 4-glucose is abundant in the CW of *S. indica.* CW: cell wall; EPS: extracellular polysaccharide; LysM: lysine motif; *p*: pyranose; *Si*: *Serendipita indica,* WSC: cell wall integrity and stress response component.

### *S. indica* EPS matrix and CW have different sugar compositions

The differential enrichment of glycan-binding proteins within the EPS matrix and CW layer prompted us to investigate the glycan composition of these two compartments isolated from *S. indica* grown in TSB medium. Protein-free EPS matrix and CW preparations were subjected to glycosyl linkage analysis for neutral sugars (Ciucanu, 2006; Liu *et al.*, 2015). About 85% of the detected glycosidic linkages could be annotated based on the retention times and the mass spectra profile of the sugar residues (Table S3 and S4). The relative abundance of 3-glucose and 3,6-glucose was higher (approx. 35%) in the EPS matrix compared to other glycosidic sugar residues (Figure 2C, Figure S4 and Table S3). In contrast, in the wall 4-linked glucose was more abundant (approx. 45%, Figure 2C, Figure S4 and Table S4) which suggests the presence of cello-oligosaccharides (Johnson *et al.*, 2018). This demonstrates that the EPS matrix and the CW of *S. indica* display major differences in the linkage compositions of their neutral sugars.

Next, we treated the EPS matrix and CW with β-glucanases from *T. harzianum* (TLE) and *H. pomatia* and analysed the digested products by TLC and MALDI-TOF (Figure 3A). Several glucan fragments with various degrees of polymerization could be detected in the digested fraction confirming that β-glucans are present in both layers (Figure 3B and Figure S5).

**Figure 3.**
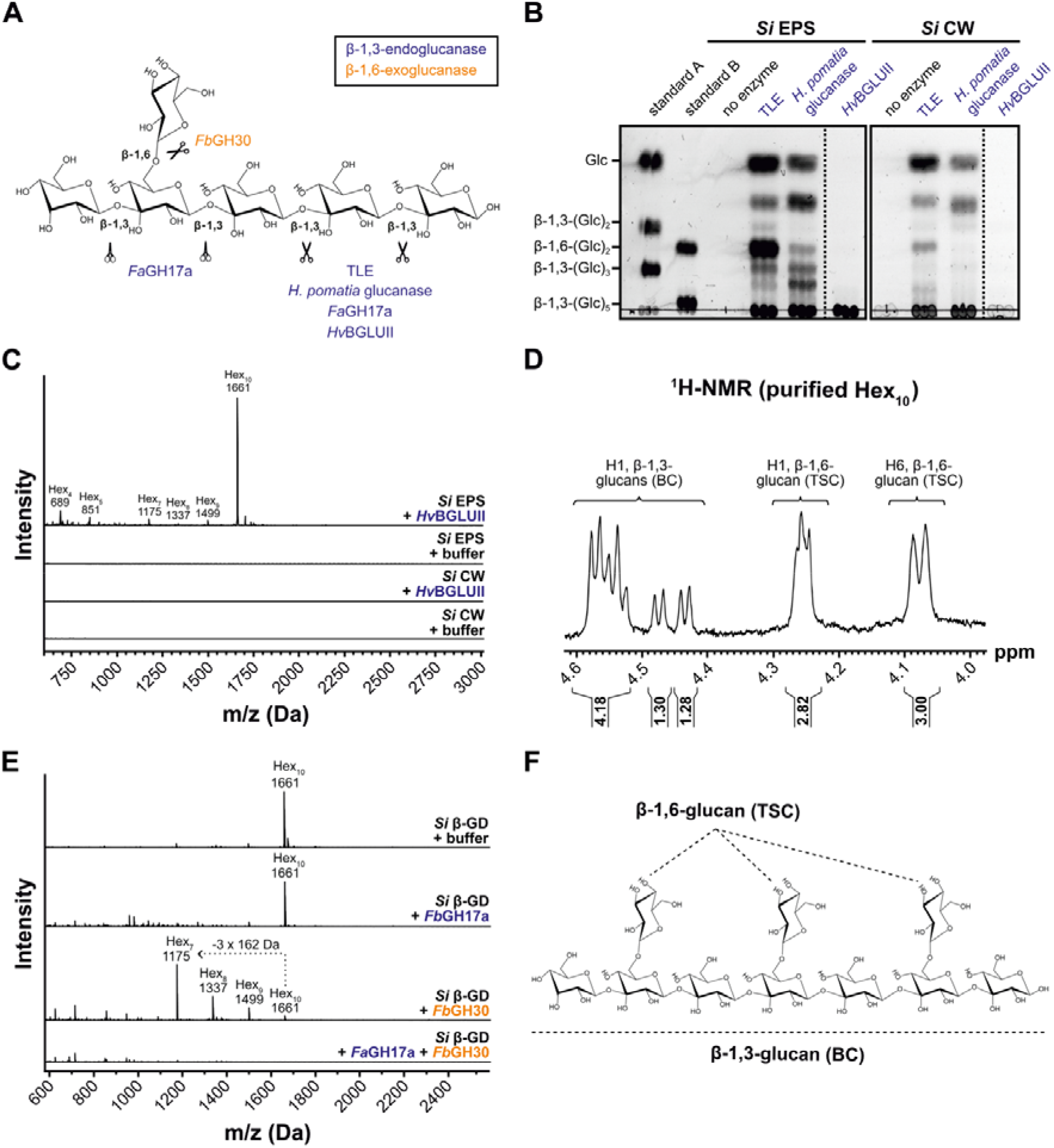
The β-1,3;1,6-glucan decasaccharide β-GD is released from the *S. indica* EPS matrix upon treatment with the barley apoplastic glycosyl hydrolase *Hv*BGLUII. (A) Glycosyl hydrolases specific for β-1,3;1,6-glucans were used for the characterization of the EPS matrix, CW and β-GD. The β-1,3-endoglucanases from *T. harzianum* (TLE) and *H. pomatia* as well as *Fa*GH17a and *Hv*BGLUII are shown as open scissors (in blue). *Fa*GH17a is represented as closed scissors because it does not hydrolyze glycosidic bonds of β-1,3-glucosyl residues substituted with β-1,6-glucosyl residues (in blue). *Fb*GH30 is a β-1,6-exoglucanase (in orange). (B) Analysis of digested EPS matrix or CW fractions by TLC. Several glucan fragments with different lengths are released from the EPS matrix and CW by the action of TLE and *H. pomatia* β-1,3-glucanase. *Hv*BGLUII releases a glucan fraction from the β-glucan-containing EPS matrix but not from the CW. (C) Analysis of digested EPS or CW fractions by MALDI-TOF mass spectrometry. The 1661 Da β-GD corresponding to 10 hexoses is released from the EPS matrix but not from the CW of *S. indica*. The representative DP of hexoses is indicated on top of the m/z (M+Na)^+^ masses of oligosaccharides. (D) ^1^H NMR spectrum of HPLC purified β-GD. (E) Treatment of β-GD with various hydrolases followed by MALDI-TOF analysis of the products. The loss of three hexoses (−3×162 Da) as a result of treatment with *Fb*GH30 is indicated with a dotted arrow. (F) Structure of the β-GD based on the ^1^H NMR spectrum. β-GD consists of a linear β-1,3-glucan backbone substituted with β-1,6-glucosyl moieties. *Si*: *Serendipita indica*; β-GD: β-1,3;1,6-glucan decasaccharide; DP: degree of polymerization; EPS: extracellular polysaccharide; Hex_n_: oligosaccharides with the indicated hexose composition; BC: backbone chain; TSC: terminal side chain.

### A β-glucan decasaccharide is released from the *S. indica* EPS matrix by a barley glucanase

We recently reported that several β-glucanases belonging to the GH17 family accumulate in the apoplast of barley roots during colonization by *S. indica* (Wawra et al., 2016). Among them, the β-glucanase *Hv*BGLUII (P15737) was consistently found at different colonization stages but also in mock-treated plants (Wawra *et al.*, 2016) (Table S5), suggesting that this may be an ubiquitous apoplastic enzyme in root tissues. To investigate the activity of *Hv*BGLUII on the fungal CW and/or on the EPS matrix, we analysed the digested fraction by TLC analysis after enzymatic incubation. Treatment of the EPS matrix with *Hv*BGLUII led to the release of a glucan fraction found at the sample origin spot that could not be further resolved under the TLC separation conditions used (Figure 3B). The corresponding band was not detected in the CW digestion (Figure 3B), indicating that *Hv*BGLUII is only active on the EPS matrix. Other β-1,3-glucanases (TLE and *H. pomatia* β-1,3-glucanase) were able to release oligosaccharides from both EPS and CW preparations, indicating that both preparations contain β-glucans that are enzyme accessible (Figure 3B and Figure S5).

To further characterize the structure of the compound released by *Hv*BGLUII from the EPS matrix, we performed MALDI-TOF and glycosyl linkage analyses. An oligosaccharide with a m/z of 1661 Da, corresponding to 10 hexoses [referred to as β-glucan decasaccharide (β-GD) fragment] with 3- and 3,6-linked glucoses was detected in high abundance compared to other oligosaccharides with various degrees of polymerization (DP4-DP9, Figure 3C and Figure S6A). Neither the β-GD nor the other oligosaccharides were detected in the supernatant of the digested CW preparation (Figure 3C), confirming the TLC result (Figure 3B). Furthermore, the β-GD was also released from the EPS matrix of *S. indica* grown in CM medium (Figure S6B), suggesting that the growth conditions do not notably influence the release of the β-GD.

To further asses the structure of β-GD, the fragment was purified using reverse-phase chromatography (Figure S7) and subjected to ^1^H NMR spectroscopy (Figure 3D). ^1^H NMR analysis displayed characteristic proton signals for β-1,3;1,6-glucan (Kim *et al.*, 2000; Lowman *et al.*, 2011; Tada *et al.*, 2009). Since β-GD was reduced prior to the NMR analysis, only 9 anomeric carbohydrate signals were present. The anomeric ^1^H NMR signals of the six internal β-1,3-glucan backbone moieties were identified at 4.4 - 4.6 ppm (Figure 3D). They appear as a multiplet due to overlapping proton doublet signals at 4.5 - 4.6 ppm (4 protons), a doublet at 4.48 ppm (J 7.8 Hz, 1 proton) representing the second glucose unit next to the reducing end and a doublet at 4.44 ppm (J 7.8 Hz, 1 proton) representing the non-reducing end of the oligosaccharide backbone. Anomeric NMR signals were also observed for the β-1,6-D-side-chain substituents at 4.26 ppm (3 protons), indicating that the oligosaccharide contains three individual monomeric substituents. This is confirmed by the H6 NMR signal of β-1,6-D-glucose substituents at 4.08 ppm (3 protons). Taken together, the ^1^H NMR analysis indicates that β-GD consists of seven β-1,3-linked D-glucose backbone units substituted with three terminal β-1,6-glucose units. The order of the substituents on the backbone could not be established by the NMR analysis performed here.

To validate the ^1^H NMR results, we took advantage of the two well-characterized glycosyl hydrolases, *Fa*GH17a and *Fb*GH30 (Becker *et al.*, 2017; Wanke *et al.*, 2020). *Fa*GH17a is an endoglucanase specifically active on unsubstituted β-1,3-glucans and *Fb*GH30 is an exoglucanase specific for β-1,6-glycosidic linkages (Figure 3A). The β-GD was treated with *Fa*GH17a or *Fb*GH30 or a combination of the two enzymes and the digested samples were analyzed using MALDI-TOF mass spectrometry (Figure 3E). Digestion with *Fb*GH30 resulted in ion signals that represent the enzymatic removal of one (m/z 1499), two (m/z 1337) or most pronounced three glucosyl moieties (m/z 1175), confirming the presence of three β-1,6-glucose units in β-GD. Digestion with *Fa*GH17a alone did not alter the molecular weight of β-GD indicating that potential enzyme hydrolysis sites are blocked by its side chain substituents. In contrast, the combined treatment with both enzymes led to complete hydrolysis of the β-GD (Figure 3E). Taken together, these results demonstrate that the β-GD released from the EPS matrix by the action of *Hv*BGLUII is a decasaccharide with seven β-1,3-glucosyl units substituted with three β-1,6-glucosyl units (Figure 3F).

### The apoplastic *Hv*BGLUII fosters MAMP-triggered immunity which is counteracted by the uncleavable β-GD

Since β-glucans represent an important class of microbial cell surface glycans able to trigger plant immune responses (Fesel and Zuccaro, 2016; Wanke *et al.*, 2020) we performed ROS burst assays with *S. indica* CW and EPS matrix as well as with the enzymatically-released β-GD to test their immunogenic potential on barley roots. Whereas the application of the fungal MAMP chitohexaose (positive control) triggered apoplastic ROS production, incubation with *S. indica* CW and EPS matrix only marginally induced ROS production (Figure S8). Remarkably, applications of the β-GD led to significantly lower ROS levels compared to the mock treatment (Figure 4A). This prompted us to test the ability of this fragment to affect ROS levels during elicitation with different MAMPs (Figure 4A, Figure S9 and Figure S10). The combined application of chitohexaose and β-GD led to a decreased accumulation of ROS with increasing concentrations of added β-GD (Figure 4A and Figure S9). Combined digestion of the β-GD with the endoglucanases *Fa*GH17a and *Fb*GH30 restored the chitohexaose-triggered ROS burst (Figure 4B), thereby highlighting that the decreased accumulation of apoplastic ROS is linked to the presence of an - at least largely - intact β-GD. ROS suppression was not observed with *S. indica* CW or EPS matrix preparations (Figure S8). Additionally, the application of the β-GD significantly reduced ROS bursts in barley roots treated with the β-1,3;1,6-glucan laminarin and in Arabidopsis seedlings treated with the flagellin-derived peptide flg22 (Figure S10), pointing out that this effect is a universal feature of the β-GD irrespective of plant species or elicitor.

**Figure 4.**
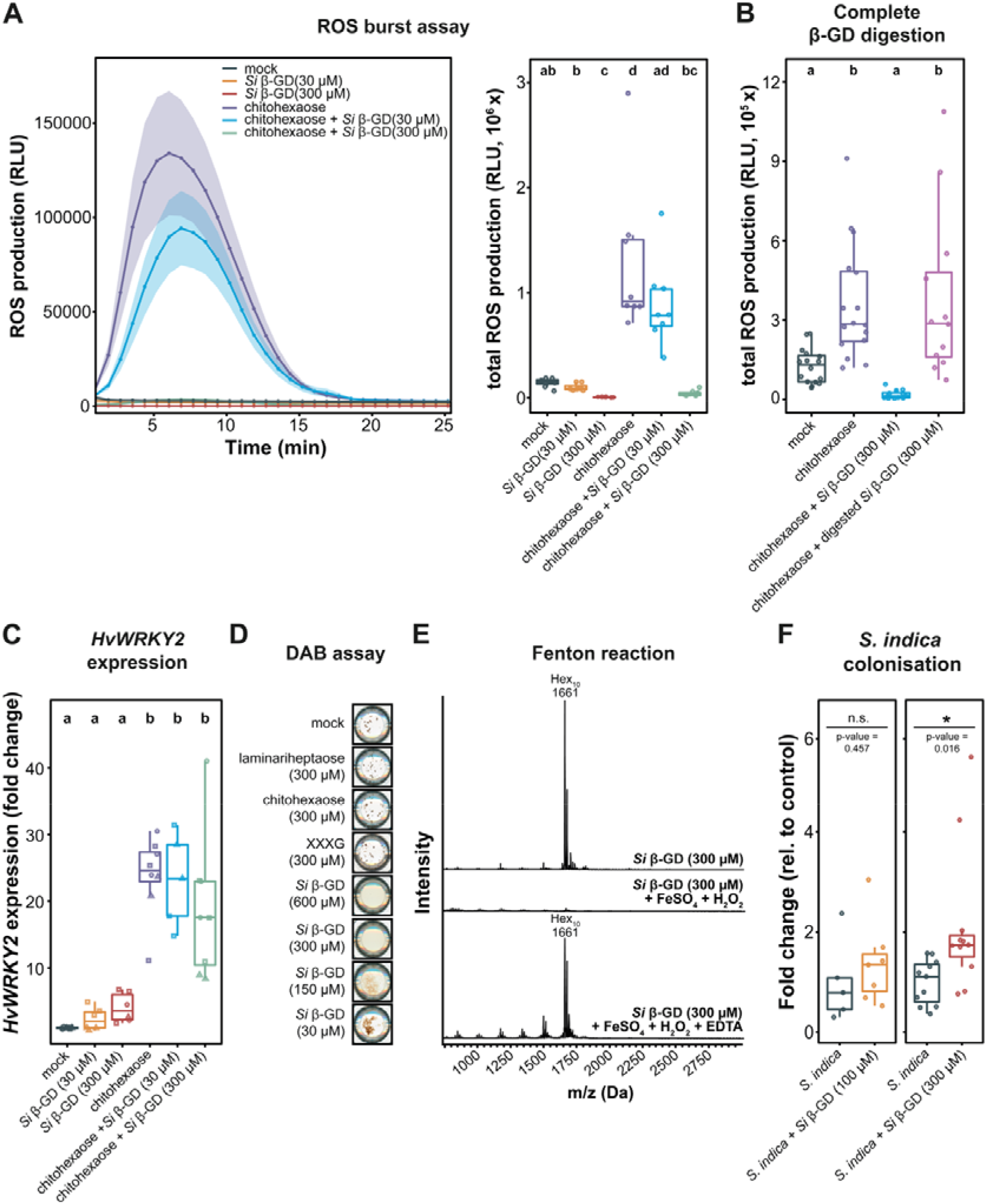
The β-GD released from *S. indica* EPS matrix scavenges ROS and enhances host colonization. (A) Apoplastic ROS production after treatment of eight-day-old barley roots with 25 μM chitohexaose and/or purified β-GD from *S. indica.* ROS production was monitored *via* a luminol-based chemiluminescence assay. Treatment with Milli-Q water was used as mock control. Boxplot represents total cumulative ROS production over the measured time period. Values represent means ± SEM from eight wells, each containing four root pieces. In total, roots from 16 individual barley plants were used per experiment. The assay was performed at least four times with independent β-GD preparations. Letters represent statistically significant differences in expression based on a one-way ANOVA and Tukey’s post-hoc test (significance threshold: p-value ≤ 0.05). (B) Before treatment of barley root pieces with the elicitors, β-GD was digested overnight (25°C, 500 rpm in heat block) with the glucanases *Fa*GH17a and *Fb*GH30, which led to complete digestion of β-GD (see also Figure 3E). As control, β-GD without the addition of enzymes (but instead with the same volume of Milli-Q water) was treated similarly. Barley root pieces were treated with Milli-Q water (n=16) and 25 μM chitohexaose alone (n=16) or in combination with digested or undigested β-GD (300 μM, n=12). Statistically significant differences are indicated by different letters based on a one-way ANOVA and Tukey’s post-hoc test (significance threshold: p-value ≤ 0.05). (C) Barley root pieces were collected one hour after elicitor treatment and further processed for RNA extraction and cDNA synthesis. Expression changes of the elicitor-responsive gene *HvWRKY2* were analysed by RT-qPCR. Fold change expression were calculated by normalization to housekeeping gene expression (*HvUBI)* and mock treatment. Data from three independent experiments are indicated by different dot shapes. Letters represent statistically significant differences in expression based on two-way ANOVA (additive model, treatment + experiment) and Tukey’s post-hoc test (significance threshold: p-value ≤ 0.05). Significant differences were associated with different treatments (F=11.629, P=1.58×10^−6^), but not with independent experiments (F=2.227, P=0.124). (D) The capability of different carbohydrates to prevent hydrogen peroxide-based and horseradish peroxidase-catalysed oxidation and precipitation of DAB was monitored. Respective sugars (or Milli-Q water as mock control) were pre-incubated with 1 mM H_2_O_2_ and 0.05 μM horseradish peroxidase before DAB (50 μM) was added. Scans of wells from 96-well plates were performed 16 hours after DAB addition. (E) Oxidative degradation of *S. indica* EPS matrix-derived β-GD (300 μM) by H_2_O_2_ was detected with an overnight Fenton reaction (1 mM H_2_O_2_, 100 μM FeSO_4_) followed by MALDI-TOF mass spectroscopic analysis. As controls, either sugar alone or the samples supplemented with 100 μM EDTA were used. (F) Colonization of barley roots by *S. indica* upon daily application of sterile Milli-Q water (mock) or β-GD (100 or 300 μM). Fungal colonization in each biological replicate was assessed by RT-qPCR comparing the expression of the fungal housekeeping gene *SiTEF* and the plant gene *HvUBI* (n=5-11). Statistical significance was determined on the non-transformed values (before normalization to *S. indica* control treatment) using a two-tailed Student’s t-test (*: p-value ≤ 0.05). β-GD: β-1,3;1,6-glucan decasaccharide; DAB: 3,3′-diaminobenzidine; EPS: extracellular polysaccharide; *Si*: *Serendipita indica*; RLU: relative light units; ROS: reactive oxygen species; XXXG: xyloglucan heptasaccharide.

To clarify whether the β-GD interferes with the MAMP perception machinery or detoxifies ROS, we tested its effect on the downstream chitohexaose-mediated induction of the *HvWRKY2* gene. *HvWRKY2* has been demonstrated to act as a reliable marker for the onset of early immune responses among a wide range of elicitors applied to barley (Liu *et al.*, 2014; Shen *et al.*, 2007; Wanke *et al.*, 2020). Despite the reduction of the oxidative burst, chitohexaose-triggered *HvWRKY2* expression was not reduced by the application of the β-GD (Figure 4C and Figure S9C). This shows that the β-GD does not prevent MAMP perception but acts on the released ROS. Furthermore, treatment with the β-GD alone did not lead to a significant increase in *HvWRKY2* expression, supporting the notion that this β-glucan fragment does not exhibit an immunogenic activity in barley roots. This is surprising due to its structural similarity to laminarihexaose and laminarin, two potent ROS elicitors in different plant species, including barley (Wanke *et al.*, 2020). Thus, the frequency and position of β-1,6-glucose substituents may define both their immunomodulatory potential as MAMP as well as their biochemical activity as ROS scavengers. To confirm this hypothesis, we treated laminariheptaose, the β-1,3-backbone of β-GD, with *Hv*BGLUII for 1 hour and overnight and tested those preparations in ROS burst assays. Barley BGLUII was capable of digesting the laminariheptaose to glucose and laminaribiose (Figure S11). Remarkably, the activity of *Hv*BGLUII on the laminariheptaose led to higher ROS accumulation compared to incubation with laminariheptaose (Figure S11). This demonstrates that *Hv*BGLUII is a host defense enzyme that releases potent MAMPs from β-1,3-glucan polymers present on the fungal surface, playing a role in host glycan perception.

### The β-GD scavenges apoplastic ROS

Numerous studies have highlighted the capability of sugars and specifically of β-glucans to act as ROS scavengers, contributing to the intracellular antioxidant system in different eukaryotes (Benaroudj *et al.*, 2001; Boulos and Nystrom, 2017; Lei *et al.*, 2015; Nishizawa *et al.*, 2008; Peshev *et al.*, 2013; Valluru and Van den Ende, 2008). To test whether the β-GD can directly act as an antioxidant, we performed an *in vitro* 3,3’-diaminobenzidine (DAB) assay. In the presence of hydrogen peroxide and horseradish peroxidase as catalyst, DAB is oxidized and polymerizes, ultimately leading to the formation of a brown, water-insoluble precipitate (Figure 4D). At low concentrations of β-GD (30-150 μM), DAB is still oxidized but forms precipitates with lower density compared to the mock control. Assuming that the β-GD is able to scavenge ROS, this effect can be explained by a decreased rate of DAB precipitation due to lower amounts of available ROS. No precipitate at all was observed at higher concentrations of β-GD between 300 and 600 μM (Figure 4D). Remarkably, other CW-associated sugars such as chitohexaose, laminariheptaose and a xyloglucan heptasaccharide (XXXG) did not interfere with DAB precipitation (Figure 4D). Mechanistically, non-enzymatic scavenging of hydroxyl radicals by sugars is based on their oxidation that can lead to cleavage of glycosidic linkages and the formation of less reactive sugar radicals that further cross-react with themselves or other sugars (Matros *et al.*, 2015). To validate if the oxidation of the decasaccharide contributes to ROS scavenging, we performed a Fenton reaction-based assay with the β-GD followed by MALDI-TOF analysis. In the presence of the Fenton reagents, the peak at 1661 Da corresponding to the β-GD was no longer detected, suggesting that the fragment might have undergone oxidative degradation or/and have changed its chemo-physical properties by the activity of hydroxyl radicals produced by the Fenton reaction (Figure 4E). The oxidative degradation of the β-GD could be rescued in the presence of EDTA, which is chelating the catalytic iron involved in the formation of hydroxyl radicals. Complete degradation was not observed for the structurally related laminariheptaose, chitohexaose and xyloglucan heptasaccharide (Figure S12). Altogether, these results demonstrate that the activity of *Hv*BGLUII on the *S. indica* EPS matrix does not release a MAMP initiating plant defense responses, but a fragment that can detoxify apoplastic ROS possibly *via* oxidative degradation. These results highlight a hitherto undescribed function of the fungal EPS matrix as a protective layer to mitigate oxidative stresses during plant-microbe interaction.

To explore whether the β-GD can facilitate root colonization during early interaction, we performed colonization assays on barley roots in the presence of various β-GD concentrations (Figure 4F). Addition of β-GD slightly increased colonization at 100 μM and led to an on average 2.15-fold increase of fungal colonization at 300 μM after 3 dpi. This demonstrates that an EPS matrix-resident decasaccharide that is released by a plant glucanase can act as a carbohydrate-class effector enhancing fungal colonization.

### The antioxidative properties of the β-GD are conserved among pathogenic and beneficial fungi

To clarify whether the observed properties of the β-GD are conserved among fungi with different lifestyles and taxonomic positions, we performed glycosyl linkage analysis of the CW and EPS matrix of *B. sorokiniana* grown in YPD medium. As observed for *S. indica*, the relative abundance of 4-linked glucose in the *B. sorokiniana* CW was higher compared to other sugar residues (Figure 5A, Figure S13 and Table S6). In the *B. sorokiniana* EPS matrix, 3-glucose and 3,6-glucose represented only a minor fraction (approx. 10%) compared to 2,3-hexose, 2,3,4-hexose and 2,3,6-hexose (approx. 40%) and this is different from the EPS matrix of *S. indica* (Figure 5A, Figure S13 and Table S7). Still, the presence of 3-glucose and 3,6-glucose suggests that β-glucans with similar structures to *S. indica* glucans are also present in the EPS matrix of *B. sorokiniana*. The EPS matrix was treated with TLE and the digested products were analysed by MALDI-TOF. Several glucan fragments with various degrees of polymerization (DP3-DP10) could be detected in the digested fraction but not in the buffer control (Figure S14), indicating that β-glucans are indeed present in the EPS matrix.

**Figure 5.**
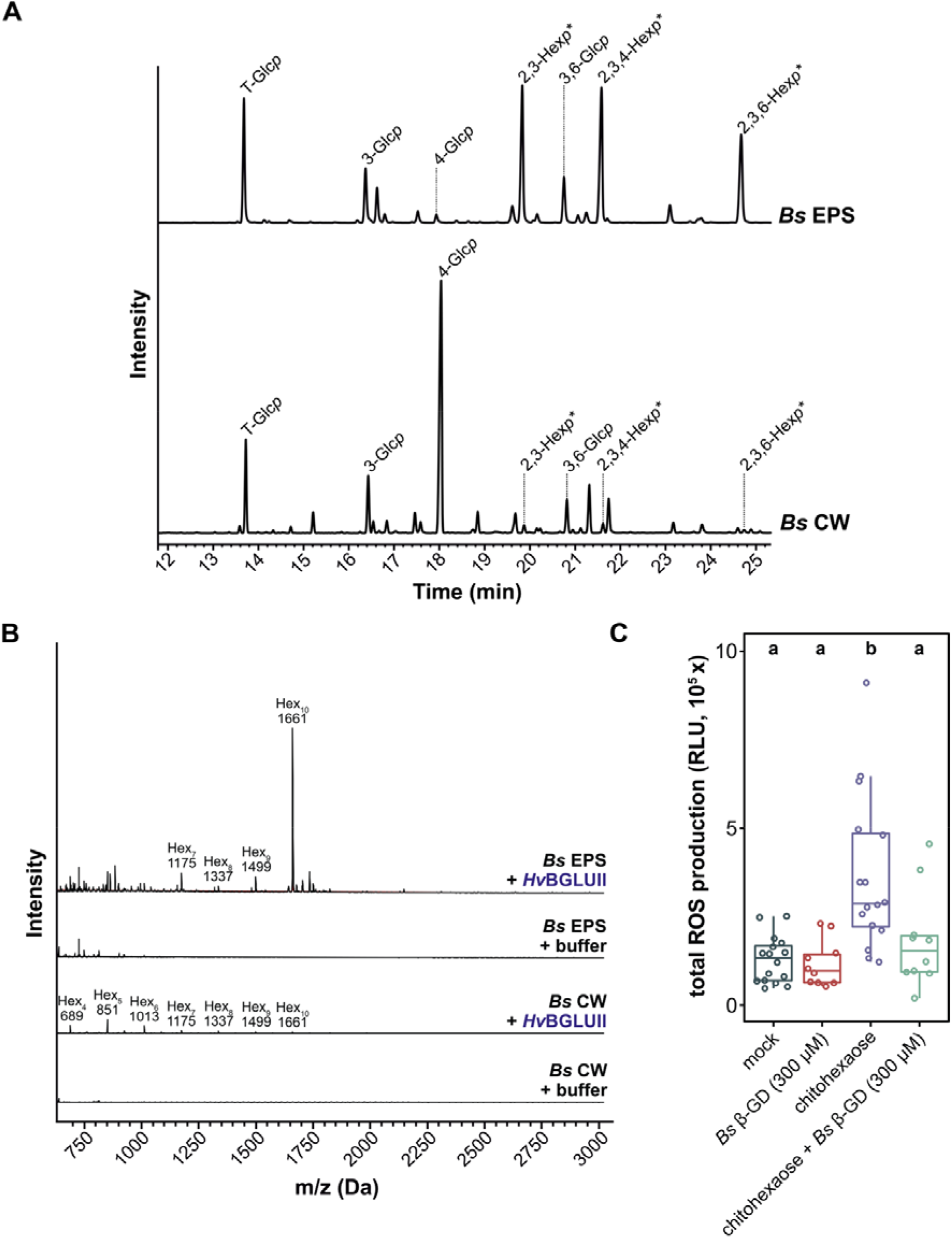
The β-GD derived from the hydrolysis of *B. sorokiniana* EPS matrix exhibits antioxidative properties. (A) Glycosidic linkage analysis of *B. sorokiniana* EPS matrix and CW preparations. 2,3-hexose, 2,3,4-hexose and 2,3,6-hexose are abundant in the EPS matrix, whereas 4-glucose is abundant in the CW of *B. sorokiniana*. (B) Analysis of digested EPS or CW fractions by MALDI-TOF mass spectrometry. The 1661 Da β-GD corresponding to 10 hexoses is released from the EPS matrix but not from the CW of *B. sorokiniana*. The representative DP of hexoses is indicated on top of the m/z (M+Na)^+^ masses of oligosaccharides. (C) ROS burst assay was performed on barley roots treated with Milli-Q water (mock), chitohexaose (25 μM), *Bs* β-GD (300 μM) or a combination of chitohexaose and *Bs* β-GD. Boxplots represent total cumulative ROS production over a measured time interval of 25 minutes. Statistically significant differences are indicated by different letters based on a one-way ANOVA and Tukey’s post-hoc test (significance threshold: p-value ≤ 0.05). β-GD: β-1,3;1,6-glucan decasaccharide; *Bs*: *Bipolaris sorokiniana*; CW: cell wall; DP: degree of polymerization; EPS: extracellular polysaccharide; *p*: pyranose, ROS: reactive oxygen species; * exact sugar moiety unknown due to the similar mass spectrum profile of hexoses (glucose, mannose or galactose).

Since *Hv*BGLUII was detected in the apoplastic fluid of *B. sorokiniana* challenged barley roots and the gene is highly induced during *B. sorokiniana* root colonization (Table S5 and Table S8), we investigated whether *Hv*BGLUII can release oligosaccharides from *B. sorokiniana* EPS matrix. Remarkably, we could detect a 1661 Da fragment in high abundance representing a decasaccharide (Figure 5B). To characterize the structure of the 1661 Da fragment from *B. sorokiniana*, we incubated it with *Fa*GH17a or *Fb*GH30 or a combination of the two enzymes. MALDI-TOF analysis revealed that this fragment has the same digestion profile and thus most likely the same structure as *S. indica* β-GD (Figure S15), demonstrating that β-GD is conserved among distantly related fungi.

Next, to investigate the immunogenic properties of *B. sorokiniana* β-GD, we tested it ROS burst assays using barley roots. Consistent with the data obtained from *S. indica*, the enzymatically-released *B. sorokiniana* β-GD, but not crude EPS matrix or CW preparations, displayed ROS scavenging activity when co-applied with chitohexaose (Figure 5C and Figure S16). Collectively, our results indicate that the antioxidative property of the β-GD is a common feature among plant-associated fungi.

## Discussion

Fungi synthetize and secrete a wide range of glycans, which are crucial determinants of microbe–microbe and microbe–host interactions. Despite variations in glycan structures and composition of the CW and the surrounding EPS matrix between fungal species, β-glucans are generally the most abundant and complex structural components. Their recognition by host receptors activates immune responses, such as accumulation of ROS and CAZymes in the extracellular apoplastic space, which hamper host colonization (El Oirdi *et al.*, 2011; Rebaque *et al.*, 2021; Wanke *et al.*, 2021; Wanke *et al.*, 2020). In order to overcome or bypass such responses, fungi employ different strategies. Common fungal countermeasures to prevent recognition and hydrolysis of their surface exposed glycans involve converting, depleting or masking highly immunoactive CW components. Although effective, these countermeasures may not always be employed because some of these glycan structures mediate important processes that are beneficial to the fungus. This explains why some glycan structures are highly conserved and cannot be extensively modified. Thus, fungi have additionally evolved apoplastic glycan-binding effector proteins that either sequester immunoactive CW-derived elicitors from the apoplast to prevent their recognition or shield them from hydrolysis. Less is known about fungal cytoplasmic effector proteins targeting the disruption of glycan signalling inside plant cells. Additionally, fungi secrete CAZyme inhibitors and proteases that cleave host hydrolytic enzymes (Ham *et al.*, 1997; Rovenich *et al.*, 2016).

Here we describe an additional common fungal counterdefensive strategy to subvert host immunity that involves the hijacking of plant apoplastic endoglucanases to release a conserved β-1,3;1,6-glucan decasaccharide with ROS scavenging properties from the extracellular polysaccharide matrix of different fungi (Figure 6). Several classes of plant proteins, called pathogenesis-related (PR) proteins, are induced in response to fungal colonization. Among these proteins the family of PR-2 proteins, which are β-1,3-endo-type glucanases, is long known (Leubner-Metzger and Meins, 1999; Stintzi *et al.*, 1993). In seed plants, β-1,3-glucanases are widely distributed, highly regulated and abundant in the apoplast or vacuoles. Besides their role in the plant response to microbial pathogens and wounding, these enzymes are also implicated in diverse physiological and developmental processes in the uninfected plant. The GH17 family member BGLUII is present in the apoplast of uninfected and infected barley roots and highly induced upon challenge with the pathogen *B. sorokiniana*. The activity of BGLUII on the fungal EPS matrix indicates that its substrate is available/exposed in this layer, the first physical site of contact with the plant defense components, but not in the CWs of two unrelated fungi. The released β-GD is not immunoactive in barley and Arabidopsis and has some specific properties, which include resilience to further digestion by GH17 family members as well as ROS scavenging abilities. These properties seem to depend on the presence of β-1,6-linked glucosyl substituents. In fact, the structurally related laminariheptaose, which has the same structure as the backbone of β-GD, is highly sensitive to digestion by *Hv*BGLUII and other GH17 family members, is immunoactive in barley, is not significantly degraded in the Fenton reaction and does not display ROS scavenging capabilities (Figure 4D and Figure S11 and S12). The removal of the side branches from the β-GD by a microbial-derived β-1,6-glucanase makes it sensitive to further digestion by GH17 family members. *In planta*, the β-GD most likely corresponds to a host glucanase-resistant structure because plants are not known to produce β-1,6-glucanases (Fliegmann *et al.*, 2004; Nars *et al.*, 2013).

**Figure 6.**
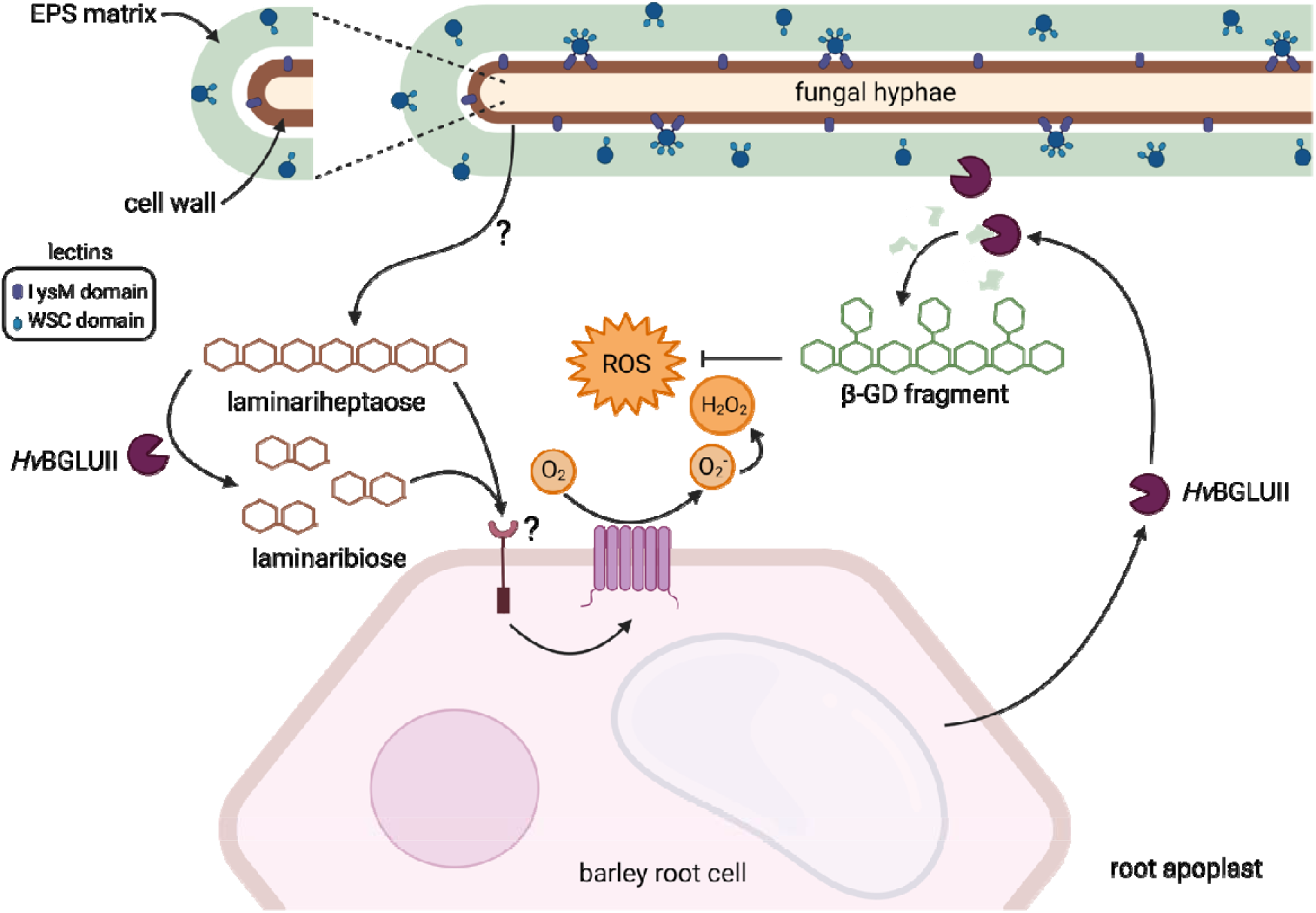
Model for the production and function of the fungal EPS-derived β-1,3;1,6-glucan decasaccharide. The fungal-responsive GH17 family member *Hv*BGLUII is found in the apoplast of barley roots and acts on β-1,3-glucans. The activity of *Hv*BGLUII leads to the production of a stronger MAMP upon hydrolysis of laminariheptaose into laminaribiose and glucose. This implicates that *Hv*BGLUII is a host-defense enzyme with a role in host glycan perception. To counteract the activity of GH17 family members plant colonizing fungi produce an EPS matrix which contains β-1,3;1,6-glucans. The activity of *Hv*BGLUII on this EPS matrix releases a conserved β-1,3;1,6-glucan decasaccharide (β-GD) which is resilient to further digestion due to the presence of the β-1,6 glucose branches and has ROS scavenging activity.

Additionally, the branching frequency might also be crucial for the immunomodulating activities of β-glucans. In animal systems, glucans are sensed by the well characterized receptor Dectin-1. The minimal glucan subunit structure for Dectin-1 activation is a β-1,3-glucan oligosaccharide containing a backbone with at least seven glucose subunits and a single β-1,6-linked side-chain branch at the non-reducing end (Adams *et al.*, 2008). We recently have shown that in plants β-1,3-glucan oligomers are perceived in a host species and length-dependent manner. While the monocots barley and *Brachypodium distachyon* can recognize longer (laminarin) and shorter (laminarihexaose) β-1,3-glucans with responses of varying intensity, duration and timing, the dicot *Nicotiana benthamiana* activates immunity in response to long β-1,3-glucans, whereas Arabidopsis and *Capsella rubella* perceive short β-1,3-glucans. The hydrolysis of the β-1,6 side-branches of laminarin did not affect recognition demonstrating that not the glycosidic decoration but rather the degree of polymerization plays a pivotal role in the recognition of long-chain β-glucans in plants. Our data do not unambiguously demonstrate the glucosyl position of the substituents on the heptasaccharide backbone of the β-GD, but a possible explanation to the observed differences between the immune properties of laminarin (1:10 branching) and the β-GD (1:2.3 branching) might be that in barley roots β-glucans with a higher degree of branching could stereochemically interfere with each other, leading to less binding by specific receptors. It could therefore well be that the β-GD has different immune properties depending on the plant species, but this requires further investigation. Nevertheless, here we show that upon incubation of Arabidopsis and barley with β-GD, no accumulation of ROS was detected. The immunomodulatory properties of the β-GD resemble those of fungal effector proteins, suggesting that it could be considered a carbohydrate-class effector with the ability to enhance fungal colonization.

In conclusion, our data demonstrate that the fungal EPS matrix is a source of soluble β-glucans that lead to ROS scavenging properties important for host colonization and represents a distinct outer fungal layer with well-defined protein and carbohydrate signatures. β-glucans can form complex higher order structures depending on the conformation of sugar residues, molecular weight, inter- and intra-chain hydrogen-bonding and environmental conditions - all effecting their properties. This is different to the immune perception of the MAMP chitin which is solely based on the length of the released oligomers by e.g. the activity of chitinases. In plants, β-glucans are considered “orphan MAMPs” as their direct immune receptors have so far not been unambiguously identified. One of the remaining challenges is to fully elucidate the β-glucans functions and receptors. Information on the structure, solubility, molecular weight, side chain branching frequency and conformation is of paramount importance and should be provided in studies dealing with fungal β-glucans and their effects on the plant immune system.

## Acknowledgments

The authors would like to thank Katharina Lufen, Institute for Plant Cell Biology and Biotechnology, Heinrich Heine University, Düsseldorf for excellent technical assistance and Alba Silipo Department of Chemical Sciences, University of Napoli Federico II for initial support with the sugar analyses. We thank Gereon Poschmann, Anja Stefanski, Thomas Lenz and Kai Stühler, Institute of Molecular Medicine, Proteome research facility, Medical Faculty and University Hospital Düsseldorf for excellent proteomics. NMR analysis was performed at the Center for Molecular and Structural Analytics at the Heinrich Heine University, Düsseldorf. We further thank Jan-Hendrik Hehemann for providing aliquots of *Fa*GH17a and *Fb*GH30. Graphical illustration was designed with the BioRender online tool.

## Contributions

BC, AW, SW and AZ conceived the study. SW established the protocols for the CW and EPS matrix extraction and performed the proteomics of EPS, CW and culture filtrates. BC performed carbohydrate analytics (glycosyl linkage analysis of EPS and CW, MALDI-TOF, purification of β-GD for ^1^H NMR and ROS burst assay). AW performed DAB assays and gene expression analysis. AW and BC performed β-GD oxidation assays. AW, SW, PS, MN, MM, MT, NC performed the ROS burst assays, EPS and CW preparations and enzymatic digestions. LM performed the colonization assays. MD contributed to the ^1^H NMR result interpretation. MP directed and supervised the carbohydrate analytics. AZ supervised the project, designed the experiments and wrote the paper with the contribution of all authors.

## References

Adams EL, Rice PJ, Graves B, Ensley HE, Yu H, Brown GD, Gordon S, Monteiro MA, Papp-Szabo E, Lowman DW, Power TD, Wempe MF, Williams DL. 2008. Differential high-affinity interaction of dectin-1 with natural or synthetic glucans is dependent upon primary structure and is influenced by polymer chain length and side-chain branching. The Journal of Pharmacology and Experimental Therapeutics 325, 115–123.

Becker S, Scheffel A, Polz MF, Hehemann J-H. 2017. Accurate Quantification of Laminarin in Marine Organic Matter with Enzymes from Marine Microbes. Applied and Environmental Microbiology 83.

Benaroudj N, Lee DH, Goldberg AL. 2001. Trehalose accumulation during cellular stress protects cells and cellular proteins from damage by oxygen radicals. The Journal of Biological Chemistry 276, 24261–24267.

Boulos S, Nystrom L. 2017. Complementary Sample Preparation Strategies for Analysis of Cereal beta-Glucan Oxidation Products by UPLC-MS/MS. Front Chem 5, 90.

Brown GD, Gordon S. 2005. Immune recognition of fungal β-glucans. Cellular Microbiology 7, 471–479.

Ciucanu I. 2006. Per-O-methylation reaction for structural analysis of carbohydrates by mass spectrometry. Analytica Chimica Acta 576, 147–155.

El Oirdi M, El Rahman TA, Rigano L, El Hadrami A, Rodriguez MC, Daayf F, Vojnov A, Bouarab K. 2011. Botrytis cinerea manipulates the antagonistic effects between immune pathways to promote disease development in tomato. Plant Cell 23, 2405–2421.

Fesel PH, Zuccaro A. 2016. β-glucan: Crucial component of the fungal cell wall and elusive MAMP in plants. Fungal genetics and biology: FG & B 90, 53–60.

Fliegmann J, Mithofer A, Wanner G, Ebel J. 2004. An ancient enzyme domain hidden in the putative beta-glucan elicitor receptor of soybean may play an active part in the perception of pathogen-associated molecular patterns during broad host resistance. The Journal of Biological Chemistry 279, 1132–1140.

Goodridge HS, Wolf AJ, Underhill DM. 2009. Beta-glucan recognition by the innate immune system. Immunological Reviews 230, 38–50.

Gow NAR, Latge JP, Munro CA. 2017. The Fungal Cell Wall: Structure, Biosynthesis, and Function. Microbiol Spectr 5.

Günl M, Kraemer F, Pauly M. 2011. Oligosaccharide mass profiling (OLIMP) of cell wall polysaccharides by MALDI-TOF/MS. Methods in Molecular Biology (Clifton, N.J.) 715, 43–54.

Ham K-S, Wu S-C, Darvill AG, Albersheim P. 1997. Fungal pathogens secrete an inhibitor protein that distinguishes isoforms of plant pathogenesis-related endo-β-1,3-glucanases. The Plant Journal 11, 169–179.

Han B, Baruah K, Cox E, Vanrompay D, Bossier P. 2020. Structure-Functional Activity Relationship of β-Glucans From the Perspective of Immunomodulation: A Mini-Review. Frontiers in Immunology 11.

Johnson JM, Thurich J, Petutschnig EK, Altschmied L, Meichsner D, Sherameti I, Dindas J, Mrozinska A, Paetz C, Scholz SS, Furch ACU, Lipka V, Hedrich R, Schneider B, Svatos A, Oelmuller R. 2018. A Poly(A) Ribonuclease Controls the Cellotriose-Based Interaction between Piriformospora indica and Its Host Arabidopsis. Plant Physiol 176, 2496–2514.

Kang X, Kirui A, Muszynski A, Widanage MCD, Chen A, Azadi P, Wang P, Mentink-Vigier F, Wang T. 2018. Molecular architecture of fungal cell walls revealed by solid-state NMR. Nat Commun 9, 2747.

Kim YT, Kim EH, Cheong C, Williams DL, Kim CW, Lim ST. 2000. Structural characterization of beta-D-(1 --> 3, 1 --> 6)-linked glucans using NMR spectroscopy. Carbohydrate Research 328, 331–341.

Lahrmann U, Ding Y, Banhara A, Rath M, Hajirezaei MR, Döhlemann S, Wirén Nv, Parniske M, Zuccaro A. 2013. Host-related metabolic cues affect colonization strategies of a root endophyte. Proceedings of the National Academy of Sciences 110, 13965–13970.

Lei N, Wang M, Zhang L, Xiao S, Fei C, Wang X, Zhang K, Zheng W, Wang C, Yang R, Xue F. 2015. Effects of Low Molecular Weight Yeast beta-Glucan on Antioxidant and Immunological Activities in Mice. Int J Mol Sci 16, 21575–21590.

Leubner-Metzger G, Meins F. 1999. Functions and regulation of plant ß-1,3-glucanases (PR-2): Pathogenesis-related proteins in plants. ed. / S.K. Datta; S. Muthukrishnan. Boca Raton, Florida: CRC Press LLC, 49–76.

Liu D, Leib K, Zhao P, Kogel K-H, Langen G. 2014. Phylogenetic analysis of barley WRKY proteins and characterization of HvWRKY1 and −2 as repressors of the pathogen-inducible gene HvGER4c. Molecular Genetics and Genomics 289, 1331–1345.

Liu L, Paulitz J, Pauly M. 2015. The Presence of Fucogalactoxyloglucan and Its Synthesis in Rice Indicates Conserved Functional Importance in Plants1[OPEN]. Plant Physiology 168, 549–560.

Lowman DW, West LJ, Bearden DW, Wempe MF, Power TD, Ensley HE, Haynes K, Williams DL, Kruppa MD. 2011. New Insights into the Structure of (1→3,1→6)-β-D-Glucan Side Chains in the Candida glabrata Cell Wall. PLOS ONE 6, e27614.

Matros A, Peshev D, Peukert M, Mock H-P, Ende WVd. 2015. Sugars as hydroxyl radical scavengers: proof-of-concept by studying the fate of sucralose in Arabidopsis. The Plant Journal 82, 822–839.

Mistry J, Chuguransky S, Williams L, Qureshi M, Salazar Gustavo A, Sonnhammer ELL, Tosatto SCE, Paladin L, Raj S, Richardson LJ, Finn RD, Bateman A. 2021. Pfam: The protein families database in 2021. Nucleic Acids Research 49, D412–D419.

Nars A, Lafitte C, Chabaud M, Drouillard S, Mélida H, Danoun S, Le Costaouëc T, Rey T, Benedetti J, Bulone V, Barker DG, Bono J-J, Dumas B, Jacquet C, Heux L, Fliegmann J, Bottin A. 2013. Aphanomyces euteiches cell wall fractions containing novel glucan-chitosaccharides induce defense genes and nuclear calcium oscillations in the plant host Medicago truncatula. PLOS ONE 8, e75039.

Nishizawa A, Yabuta Y, Shigeoka S. 2008. Galactinol and raffinose constitute a novel function to protect plants from oxidative damage. Plant Physiology 147, 1251–1263.

Nizam S, Qiang X, Wawra S, Nostadt R, Getzke F, Schwanke F, Dreyer I, Langen G, Zuccaro A. 2019. Serendipita indica E5lllNT modulates extracellular nucleotide levels in the plant apoplast and affects fungal colonization. EMBO reports 20, e47430.

Nostadt R, Hilbert M, Nizam S, Rovenich H, Wawra S, Martin J, Küpper H, Mijovilovich A, Ursinus A, Langen G, Hartmann MD, Lupas AN, Zuccaro A. 2020. A secreted fungal histidine- and alanine-rich protein regulates metal ion homeostasis and oxidative stress. New Phytologist 227, 1174–1188.

Peshev D, Vergauwen R, Moglia A, Hideg É, Van den Ende W. 2013. Towards understanding vacuolar antioxidant mechanisms: a role for fructans? Journal of Experimental Botany 64, 1025–1038.

Poschmann G, Seyfarth K, Besong Agbo D, Klafki H-W, Rozman J, Wurst W, Wiltfang J, Meyer HE, Klingenspor M, Stühler K. 2014. High-fat diet induced isoform changes of the Parkinson’s disease protein DJ-1. Journal of Proteome Research 13, 2339–2351.

Rebaque D, Del Hierro I, Lopez G, Bacete L, Vilaplana F, Dallabernardina P, Pfrengle F, Jorda L, Sanchez-Vallet A, Perez R, Brunner F, Molina A, Melida H. 2021. Cell wall-derived mixed-linked beta-1,3/1,4-glucans trigger immune responses and disease resistance in plants. Plant J.

Rovenich H, Zuccaro A, Thomma BPHJ. 2016. Convergent evolution of filamentous microbes towards evasion of glycan-triggered immunity. New Phytologist 212, 896–901.

Sarkar D, Rovenich H, Jeena G, Nizam S, Tissier A, Balcke GU, Mahdi LK, Bonkowski M, Langen G, Zuccaro A. 2019. The inconspicuous gatekeeper: endophytic Serendipita vermifera acts as extended plant protection barrier in the rhizosphere. New Phytologist 224, 886–901.

Shen Q-H, Saijo Y, Mauch S, Biskup C, Bieri S, Keller B, Seki H, Ulker B, Somssich IE, Schulze-Lefert P. 2007. Nuclear activity of MLA immune receptors links isolate-specific and basal disease-resistance responses. Science (New York, N.Y.) 315, 1098–1103.

Stintzi A, Heitz T, Prasad V, Wiedemann-Merdinoglu S, Kauffmann S, Geoffroy P, Legrand M, Fritig B. 1993. Plant ‘pathogenesis-related’ proteins and their role in defense against pathogens. Biochimie 75, 687–706.

Tada R, Adachi Y, Ishibashi K-i, Ohno N. 2009. An unambiguous structural elucidation of a 1,3-beta-D-glucan obtained from liquid-cultured Grifola frondosa by solution NMR experiments. Carbohydrate Research 344, 400–404.

Taylor JW, Berbee ML. 2006. Dating divergences in the Fungal Tree of Life: review and new analyses. Mycologia 98, 838–849.

Valluru R, Van den Ende W. 2008. Plant fructans in stress environments: emerging concepts and future prospects. Journal of Experimental Botany 59, 2905–2916.

Wanke A, Malisic M, Wawra S, Zuccaro A. 2021. Unraveling the sugar code: the role of microbial extracellular glycans in plant-microbe interactions. J Exp Bot 72, 15–35.

Wanke A, Rovenich H, Schwanke F, Velte S, Becker S, Hehemann J-H, Wawra S, Zuccaro A. 2020. Plant species-specific recognition of long and short β-1,3-linked glucans is mediated by different receptor systems. The Plant Journal 102, 1142–1156.

Wawra S, Fesel P, Widmer H, Neumann U, Lahrmann U, Becker S, Hehemann J-H, Langen G, Zuccaro A. 2019. FGB1 and WSC3 are in planta-induced β-glucan-binding fungal lectins with different functions. New Phytologist 222, 1493–1506.

Wawra S, Fesel P, Widmer H, Timm M, Seibel J, Leson L, Kesseler L, Nostadt R, Hilbert M, Langen G, Zuccaro A. 2016. The fungal-specific β-glucan-binding lectin FGB1 alters cell-wall composition and suppresses glucan-triggered immunity in plants. Nature Communications 7, 13188.

Zuccaro A, Lahrmann U, Güldener U, Langen G, Pfiffi S, Biedenkopf D, Wong P, Samans B, Grimm C, Basiewicz M, Murat C, Martin F, Kogel K-H. 2011. Endophytic Life Strategies Decoded by Genome and Transcriptome Analyses of the Mutualistic Root Symbiont Piriformospora indica. PLOS Pathogens 7, e1002290.

