## Supplementary Figures S1 to S16 for "Fungi hijack a plant apoplastic endoglucanase to release a ROS scavenging β-glucan decasaccharide to subvert immune responses"

^#^Shared first

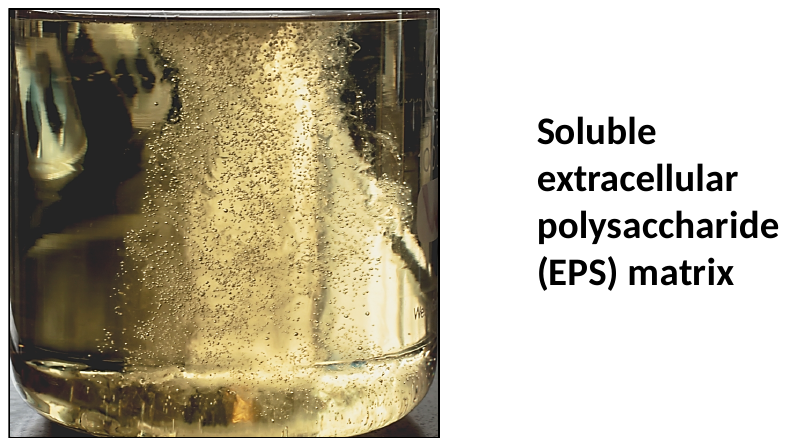

**Figure S1. EPS matrix isolated from *S. indica* using the cryogelation approach.** Air bubbles were embedded in the gel-like EPS matrix after stirring to make it visible.

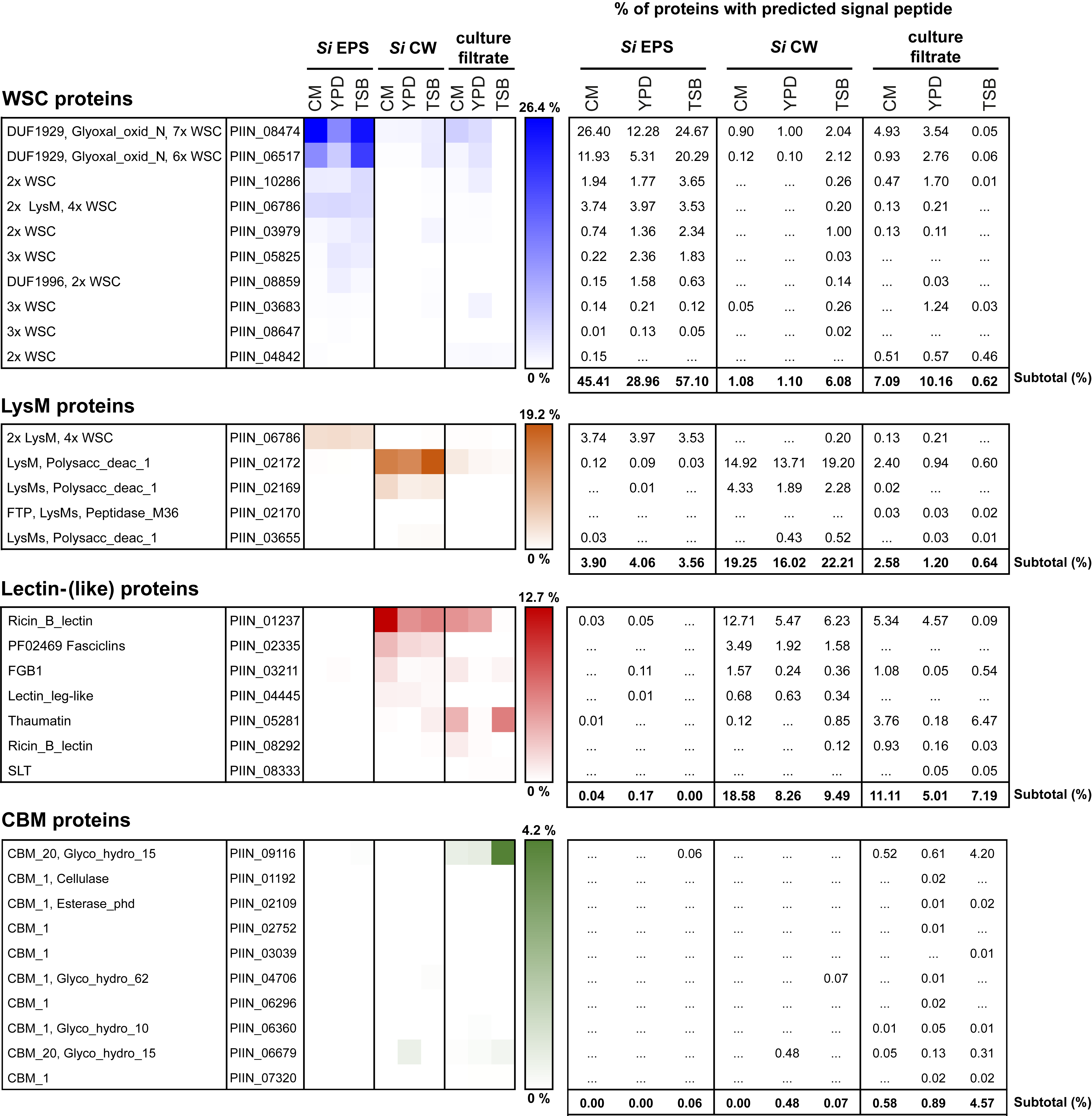

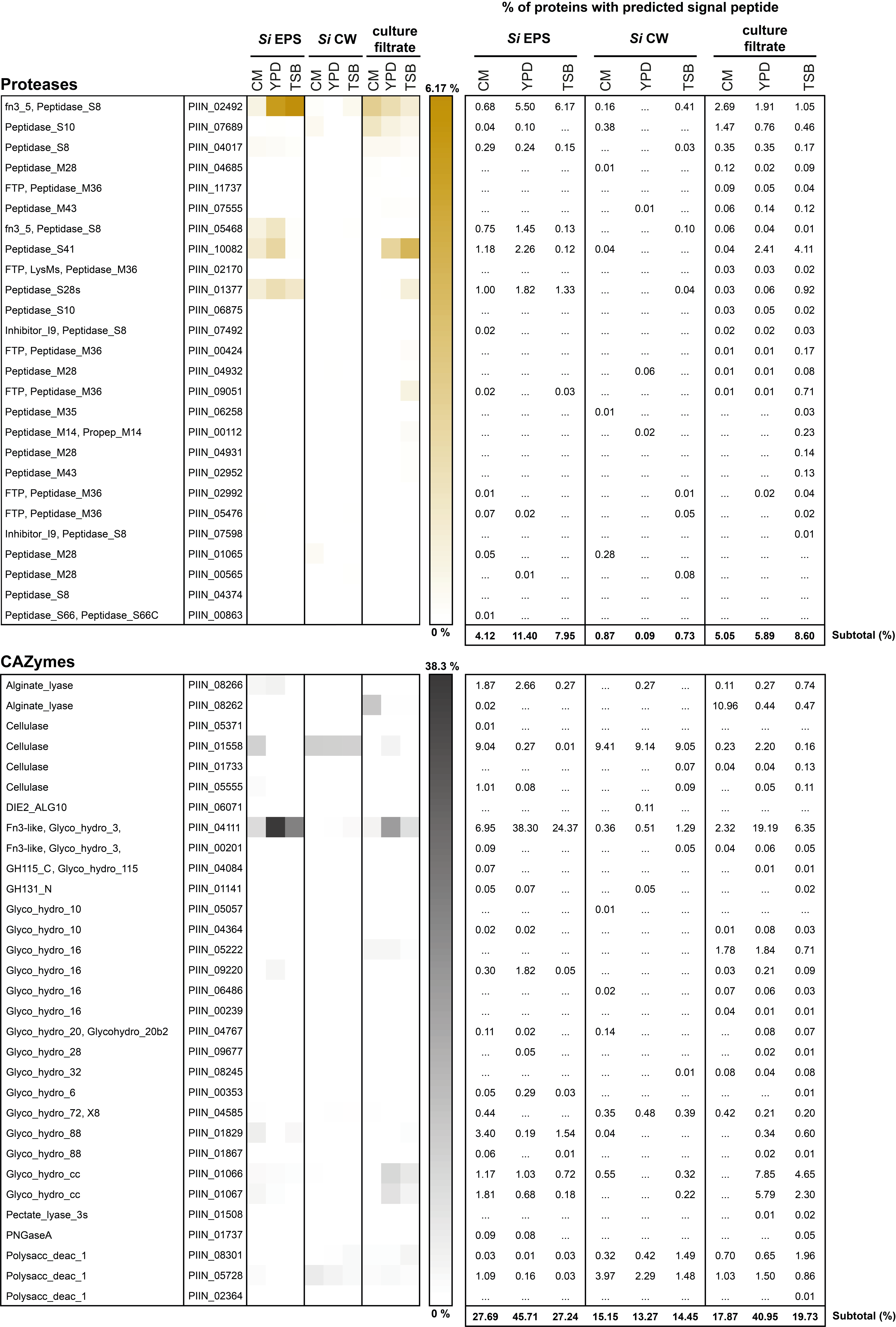

**Figure S2. Proteome analysis of *S. indica* EPS matrix, CW and culture filtrate.** *S. indica* spores were inoculated into three different liquid culture media: CM, YPD and TSB. The EPS matrix and CW proteins were isolated and solubilized by adding SDS-gel loading buffer and boiling at 95°C. The proteins present in the culture filtrate were precipitated using trichloroacetic acid and solubilized by boiling in SDS-gel loading buffer. The proteins were separated using a 10% SDS PAGE gel for 15 minutes and subsequently stained with Coomassie Brilliant Blue. The visible protein band in each sample was cut out and prepared for trypsin digest. The peptides were analyzed using LC-MS. Proteins with a predicted signal peptide were annotated using the protein families database Pfam. The relative abundance was calculated using the label-free quantification intensities values and expressed as a percentage (%). The proteome analysis of the EPS matrix and CW revealed that several WSC domain-containing proteins, including *Si*WSC3 (PIIN_05825*)* are abundant in the matrix compared to the CW. Additionally, two polysaccharide deacetylases with a LysM domain are enriched in the CW of *S. indica*. This suggests that the detected polysaccharide deacetylases with LysM domain might be involved in chitin remodeling within the CW of *S. indica.* Furthermore, a thaumatin family protein (PIIN_05281) is enriched in the culture filtrate. Thaumatins have been shown to exhibit antifungal activity through their β-1,3-glucanase activity. It is possible that PIIN_05281 exhibits an antimicrobial function against fungal competitors. Likewise, peptidases and glycosyl hydrolases with fibronectin type III module (Fn3) domains are enriched in the matrix and the culture medium but not in the CW. Fn3 are versatile domains in proteins that play a major role in mediating protein-protein interactions. CW: cell wall; EPS: extracellular polysaccharide; Fn3: fibronectin type III module; LysM: lysine motive; *Si*: *Serendipita indica*; WSC: cell wall integrity and stress response component.

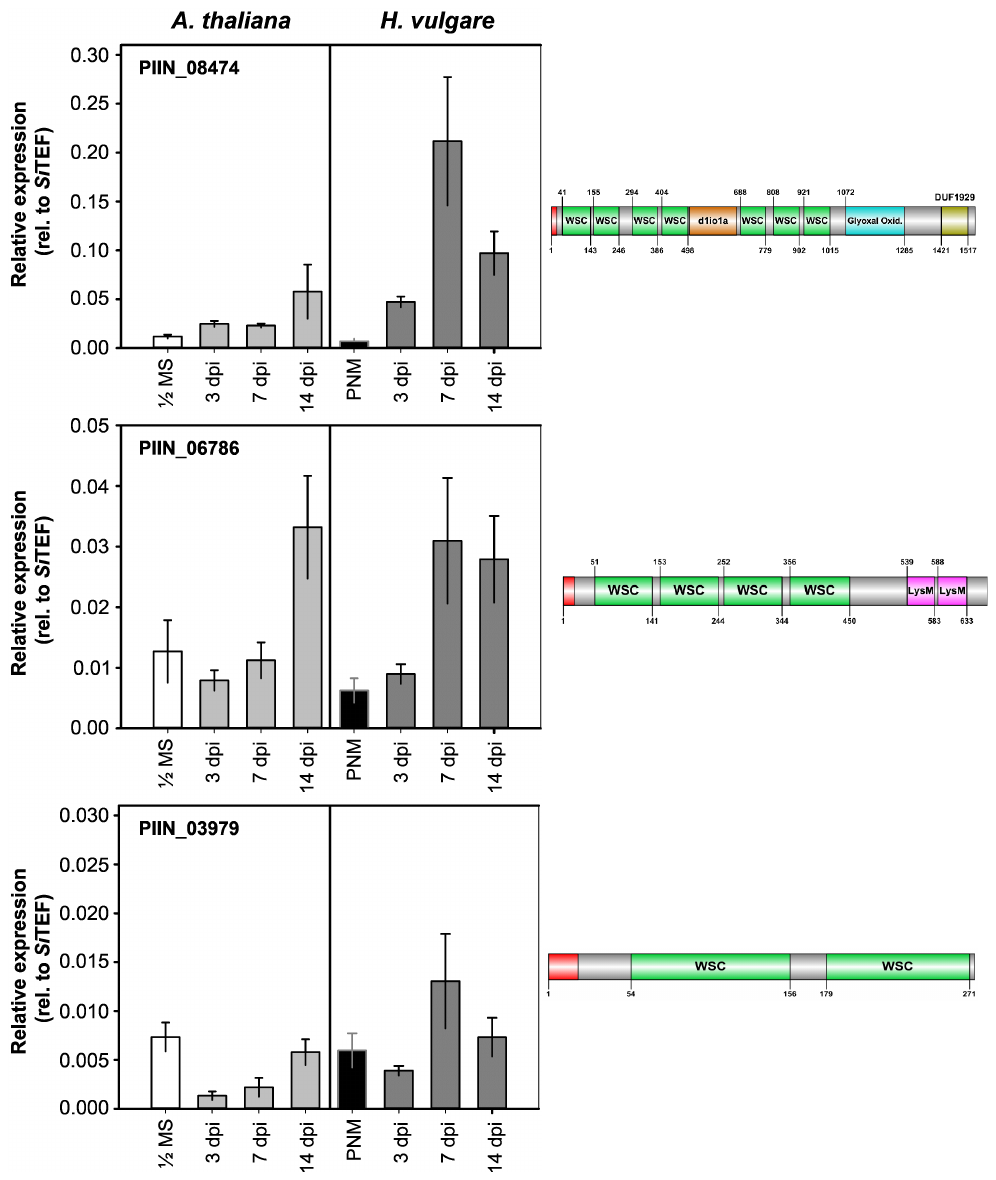

**Figure S3. Expression analysis of selected genes with WSC domains in *S. indica* during plant colonization.** The gene expression of *S. indica* WSC domain-containing proteins during root colonization of barley and Arabidopsis was monitored via quantitative reverse transcription PCR as previously described (Wawra *et al.*, 2019). Dpi: days post inoculation; LysM: lysine motive; MS: Murashige-Skoog medium; PNM: plant nutrition medium; WSC: cell wall integrity and stress response component.

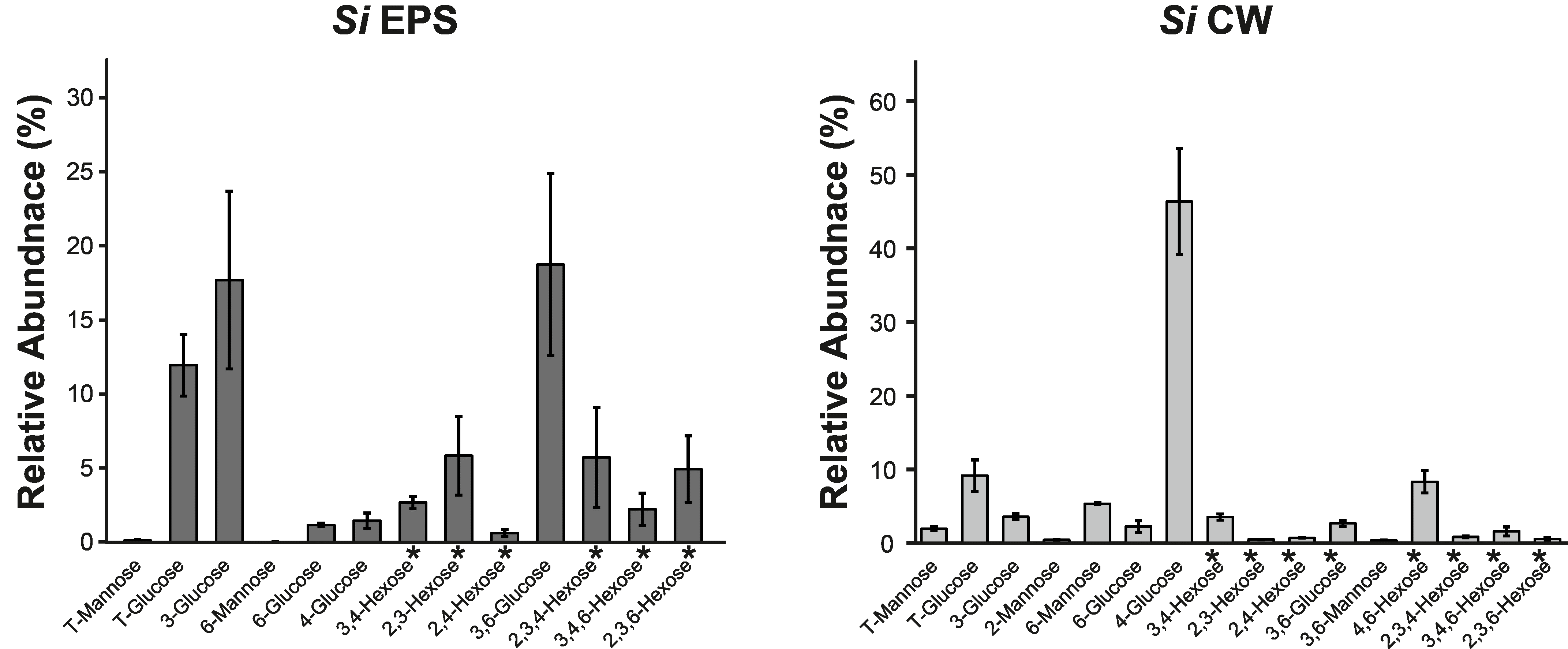

**Figure S4. Glycosyl linkage analysis of *Si* EPS and *Si* CW.** 2 mg of ground EPS matrix or the alcohol insoluble residue (CW fraction) isolated from *S. indica* grown in TSB medium containing 1% sucrose was subjected to glycosyl linkage analysis as previously described (Liu *et al.*, 2015). The sugar residues were derivatized into partially methylated alditol acetates and analyzed using GC-MS. The glycosidic linkages were assigned based on the retention time and the mass spectrum available at CCRC spectral database. The annotated glycosyl residues detected in the EPS matrix or CW are expressed as a percentage of the total glycosyl peak areas. The other minor unannotated glycosyl residues can be found in the Supplemental Table S3 and S4. Average and standard deviation of four independent biological replicates are presented. CW: cell wall; EPS: extracellular polysaccharide matrix; *Si*: *Serendipita indica;* * exact sugar moiety unknown due to the similar mass spectrum profile of hexoses (glucose, mannose or galactose).

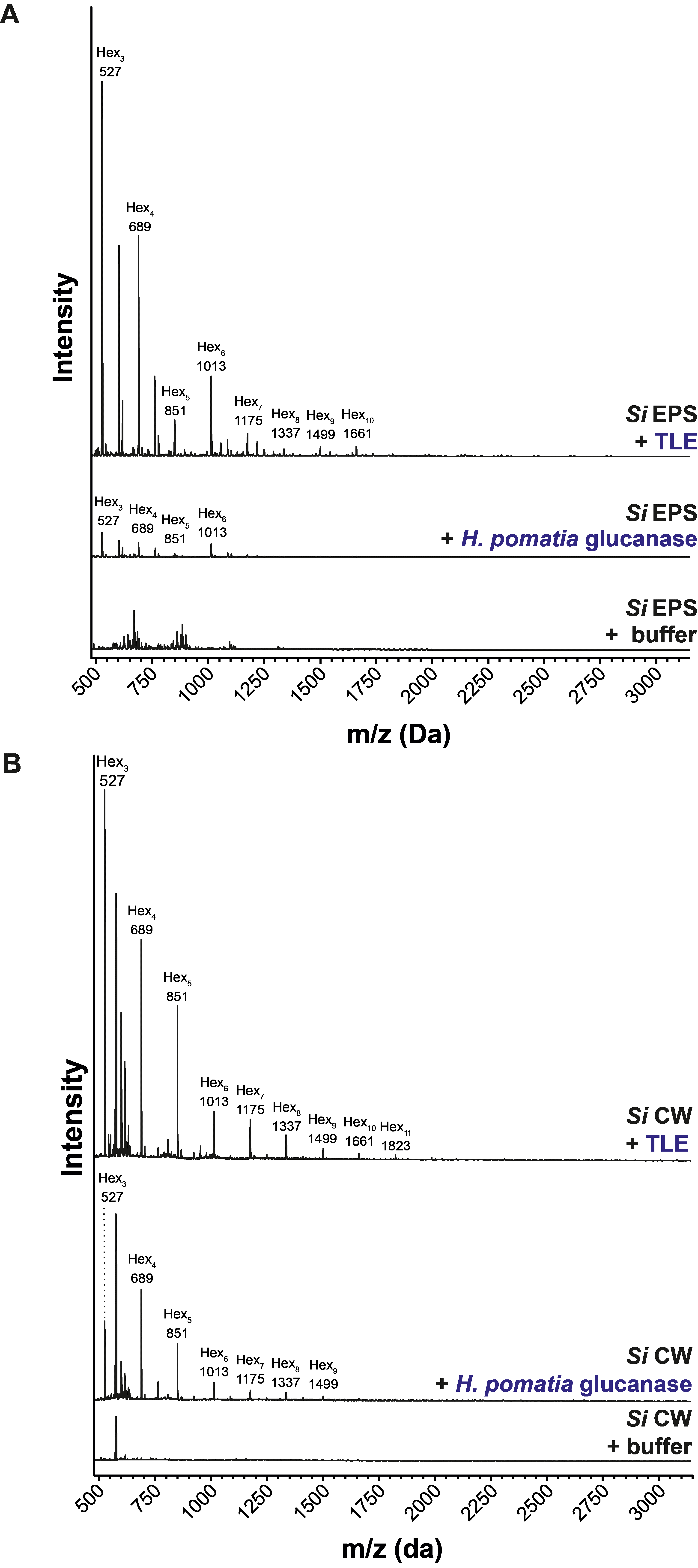

**Figure S5. Analysis of oligosaccharides released from the EPS or protein-free CW of *S. indica* by the action of β-1,3-glucanases.** 1 mg of freeze-dried EPS (A) or CW (B) obtained from *S. indica* was soaked in 2 mM sodium acetate, pH 5.0 (TLE), or 2 mM MES pH 5.0 (*H. pomatia* glucanase) at 70°C overnight. The soaked material was incubated with TLE or *H. pomatia* β-1,3-glucanase at 40°C for 16 hours. The supernatant fraction from the digested EPS matrix and CW were analyzed by MALDI-TOF. The m/z (M+Na)^+^ Da of oligosaccharides resulting from the digestion of the samples are labeled with their hexose composition. CW: cell wall; EPS: extracellular polysaccharide; Hex_n,_: oligosaccharides with the indicated hexose composition; *Si*: *Serendipita indica*; TLE: *Trichoderma* *harzianum* lysing enzymes.

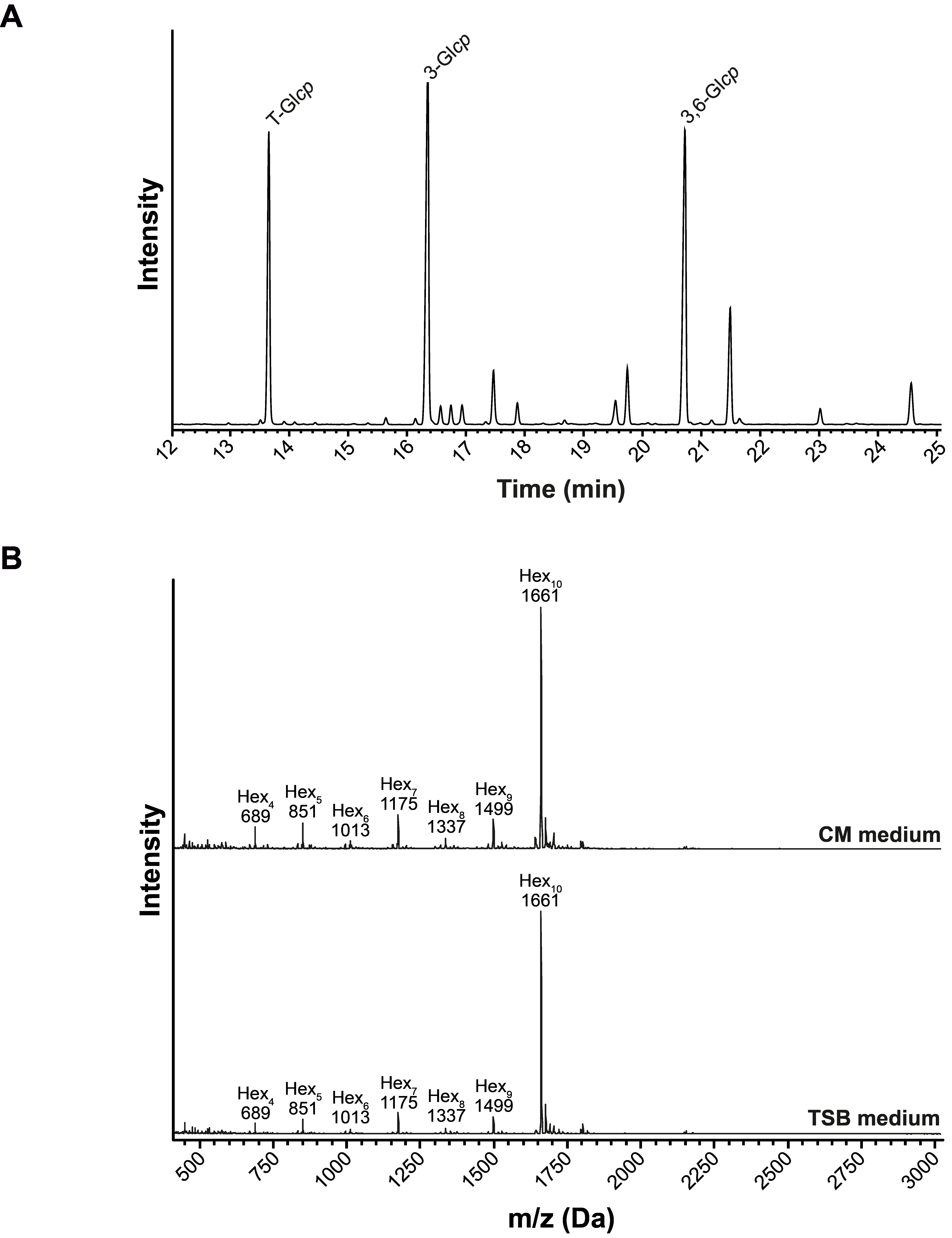

**Figure S6. Glycosyl linkage and MALDI-TOF analysis of the glucan fraction (β-GD) released from the EPS matrix of *S. indica.*** (A) Glycosyl linkage analysis of β-GD released from the EPS matrix of *S. indica*. (B) *S. indica* was grown either in liquid CM or TSB medium containing 1% sucrose. β-GD was isolated and subjected to MALDI-TOF mass spectrometry. The ions corresponding to oligosaccharides with varying degree of polymerization are labelled with their hexose composition. CM: complex Hill-Käfer medium; EPS: extracellular polysaccharide; Hex_n_: oligosaccharides with the indicated hexose composition; TSB: Tryptic Soy Broth.

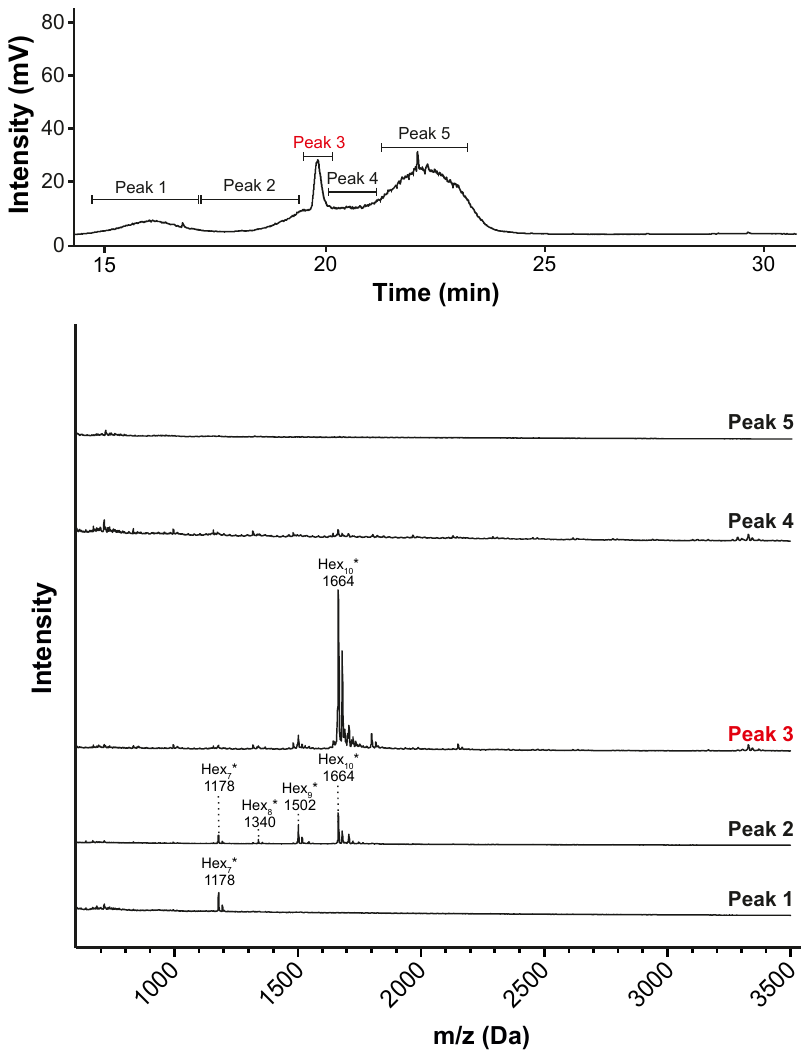

**Figure S7. Purification of the β-GD fragment for ^1^H NMR analysis.** 40 mg of the β-GD enriched fraction was reduced using NaBD_4_. The reduced fraction was neutralized with acetic acid, washed with methanol and air-dried. The dried material was dissolved in 2 ml of 6% methanol and 50 µl of the sample was subjected to liquid chromatography using a reverse-phase column (Vydac, Hesperia, CA, USA) connected to an evaporative light scattering detector. The detected peaks were collected and analyzed using MALDI-TOF. The samples were injected multiple times (24x) and peak 3 (shown in red) containing the reduced 1661 Da fragment was pooled and freeze-dried. The freeze-dried material was used for the ^1^H NMR analysis. β-GD: β-1,3;1,6-glucan decasaccharide; Hex_n_: oligosaccharides with the indicated hexose composition; * represents the increase by +3 Da in the detected oligosaccharides due to the reduction of the anomeric C1 atom.

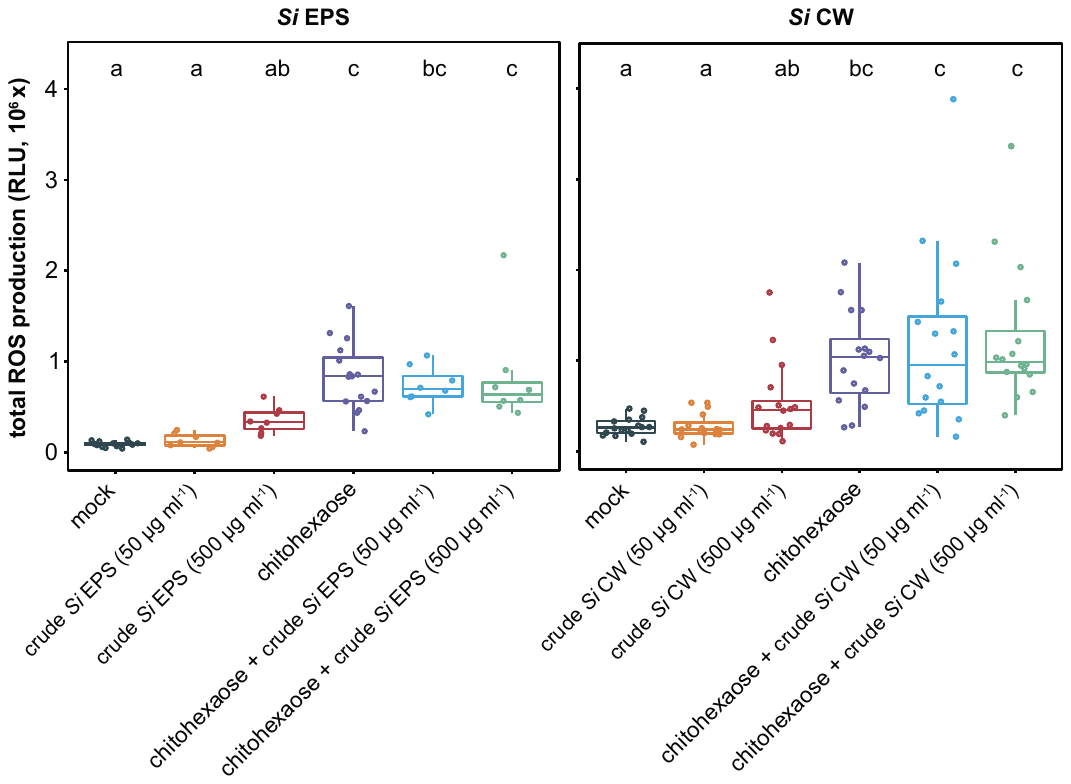

**Figure S8. Mechanically-released fragments from *S. indica* EPS matrix and CW layer do not exhibit ROS scavenging activity.** (A) *S. indica* EPS matrix and CW substrates were ground, heat-treated and sonicated prior to application to barley roots. Roots were treated with Milli-Q water (mock, n=16), chitohexaose (25 µM, n=16), crude EPS matrix or CW preparations (n=8-16) and combinations of chitohexaose and EPS matrix or CW preparations (n=8-16). Total cumulative ROS was calculated from a measured time interval of 60 minutes. Different letters indicate statistically significant differences based on a one-way ANOVA and Tukey's post‐hoc test (significance threshold: p-value ≤ 0.05). CW: cell wall; EPS: extracellular polysaccharide; RLU: relative light units; ROS: reactive oxygen species*; Si*: *Serendipita indica*.

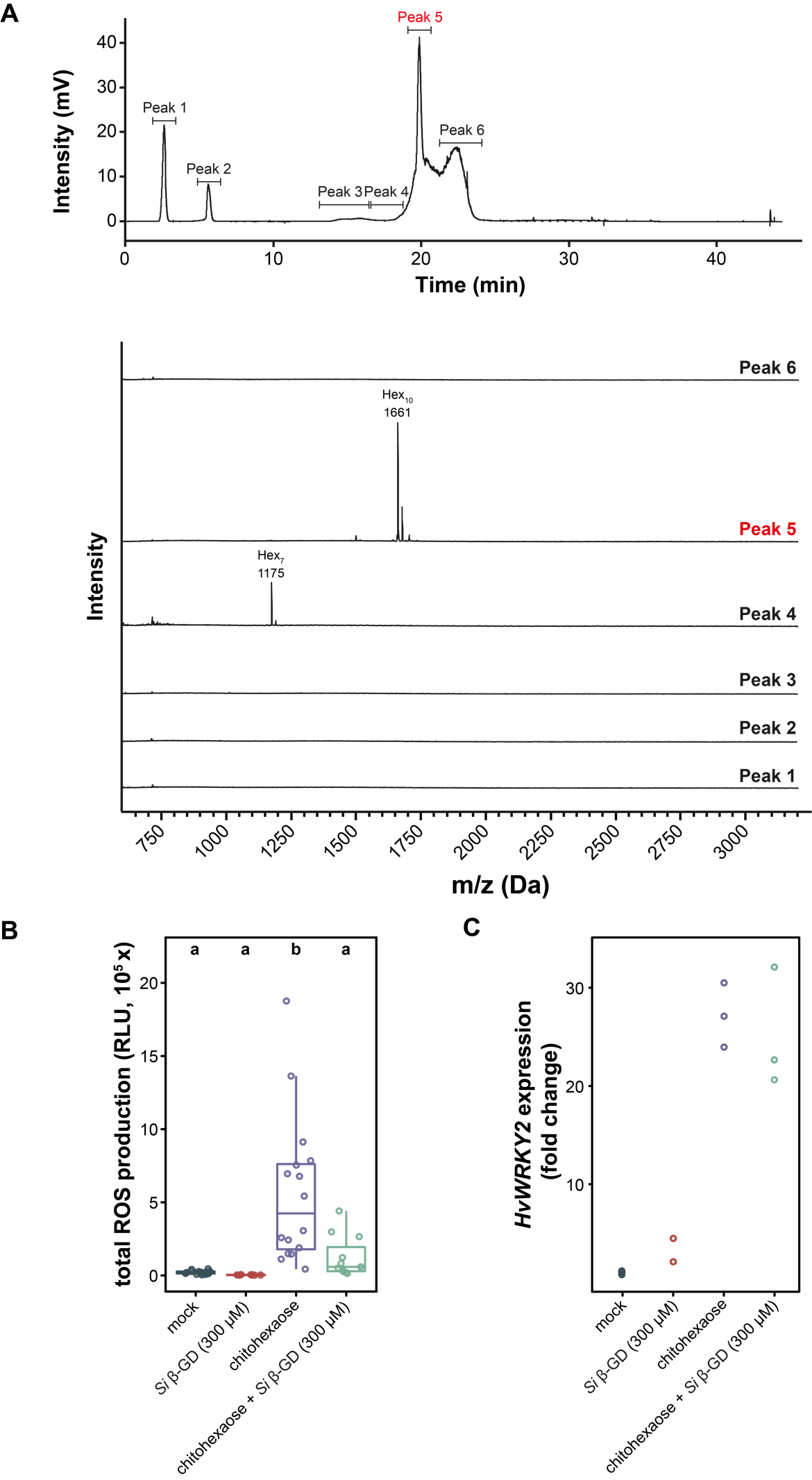

**Figure S9. Purification of native β-GD and immunogenic characterization.** (A) 40 mg of the β-GD enriched fraction was dissolved in 2 ml of 6% methanol and 50 µl of the sample was subjected to liquid chromatography using a reverse-phase column (Vydac, Hesperia, CA, USA) connected to an evaporative light scattering detector. The detected peaks were collected and analyzed using MALDI-TOF. The samples were injected multiple times (24x) and peak 5 (shown in red) containing the 1661 Da fragment was pooled and freeze-dried. HPLC-purified β-GD fraction was used in the ROS burst assay (B) and the quantitative reverse transcription PCR gene expression analysis (C) using eight-day-old barley root pieces. (B) The following treatments were applied in the ROS burst assay: mock treatment with Milli-Q water (n=16), β-GD (300 µM, n=8), chitohexaose (25 µM, n=16) and combined β-GD/chitohexaose treatments (concentrations as in previous treatments, n=11). ROS production was measured for 25 minutes. Different letters indicate statistically significant differences based on a one-way ANOVA and Tukey's post‐hoc test (significance threshold: p-value ≤ 0.05). (C) Samples for quantitative reverse transcription PCR were collected after 1 hour. Applied elicitor concentrations were as described above for the ROS burst assay (n=2-3). Gene expression of elicitor-responsive *HvWRKY2* was calculated relative to housekeeping gene expression (*HvUBI*) and normalized to mock treatment. Calculation of statistical significance was not performed due to low sample number. β-GD: β-1,3:1,6-glucan decasaccharide; RLU: relative light units; ROS: reactive oxygen species*; Si*: *Serendipita indica*.

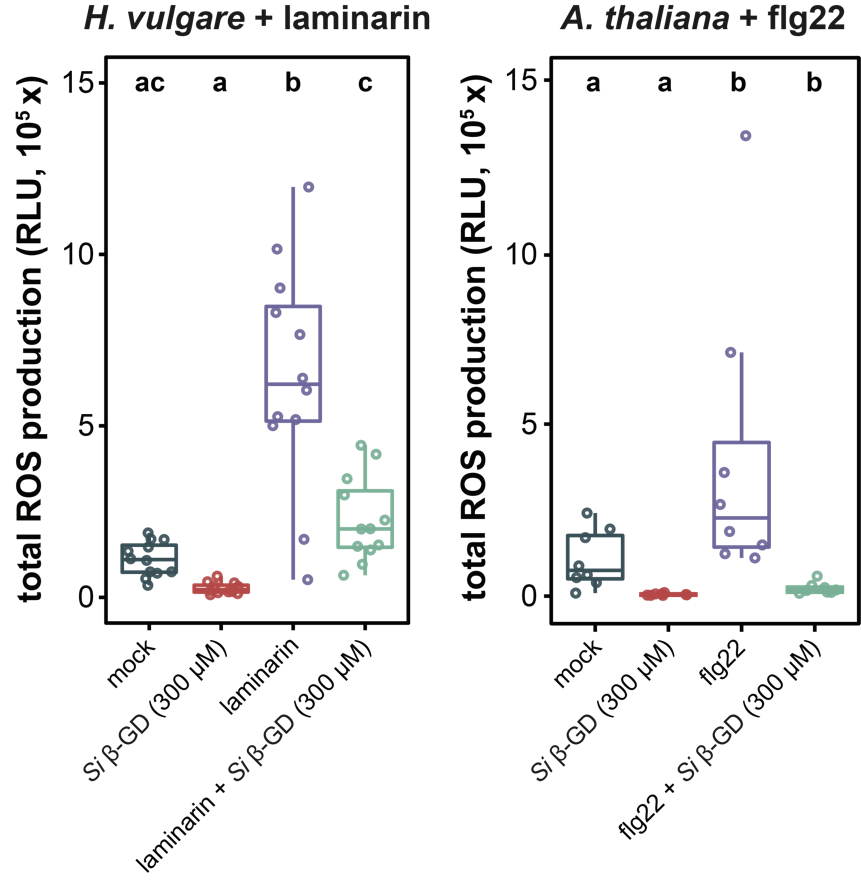

**Figure S10. Detoxification of apoplastic ROS by *S. indica* β-GD is independent of elicitor treatment and plant species.** ROS burst assays were performed with eight-day-old barley roots (n=12) and two-week-old Arabidopsis seedlings (n=8). Plants were treated with Milli-Q water (mock), 2 mg/ml laminarin,100 nM flg22, 300 µM β-GD and combinations between laminarin or flg22 and β-GD. Different letters indicate statistically significant differences based on a one-way ANOVA and Tukey's post‐hoc test (significance threshold: p-value ≤ 0.05). β-GD: β-1,3:1,6-glucan decasaccharide; RLU: relative light units; ROS: reactive oxygen species*; Si*: *Serendipita indica*.

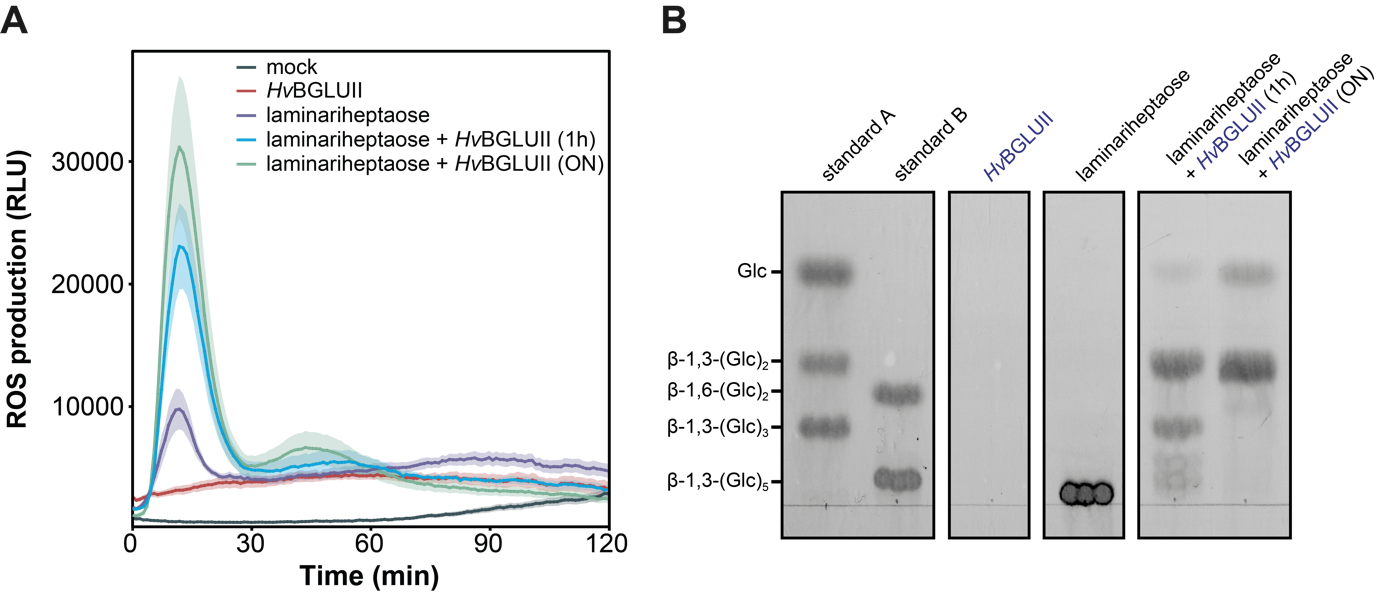

**Figure S11: Digestion of laminariheptaose with *Hv*BGLUII enhances ROS production in barley roots.** (A) Before treatment of barley roots, 3 mM laminariheptaose was digested with *Hv*BGLUII (10 units in total digestion volume of 200 µl) for one hour or overnight (ON) at 40°C, 450 rpm. As controls, Milli-Q water with *Hv*BGLUII and laminariheptaose without *Hv*BGLUII were treated in a similar way. Digestion was stopped by incubation at 90°C (10 minutes, 450 rpm) followed by centrifugation (10 min, 13000xg). Supernatants were used in ROS burst assays with eight-day-old barley roots (n=8, three root pieces per replicate). Plants were treated with Milli-Q water (mock), Milli-Q water with *Hv*BGLUII or 300 µM laminariheptaose (digested or non-digested). (B) Digestion of substrates used in ROS burst assay was analyzed via thin-layer chromatography. Glc: glucose; ON: overnight; RLU: relative light units; ROS: reactive oxygen species.

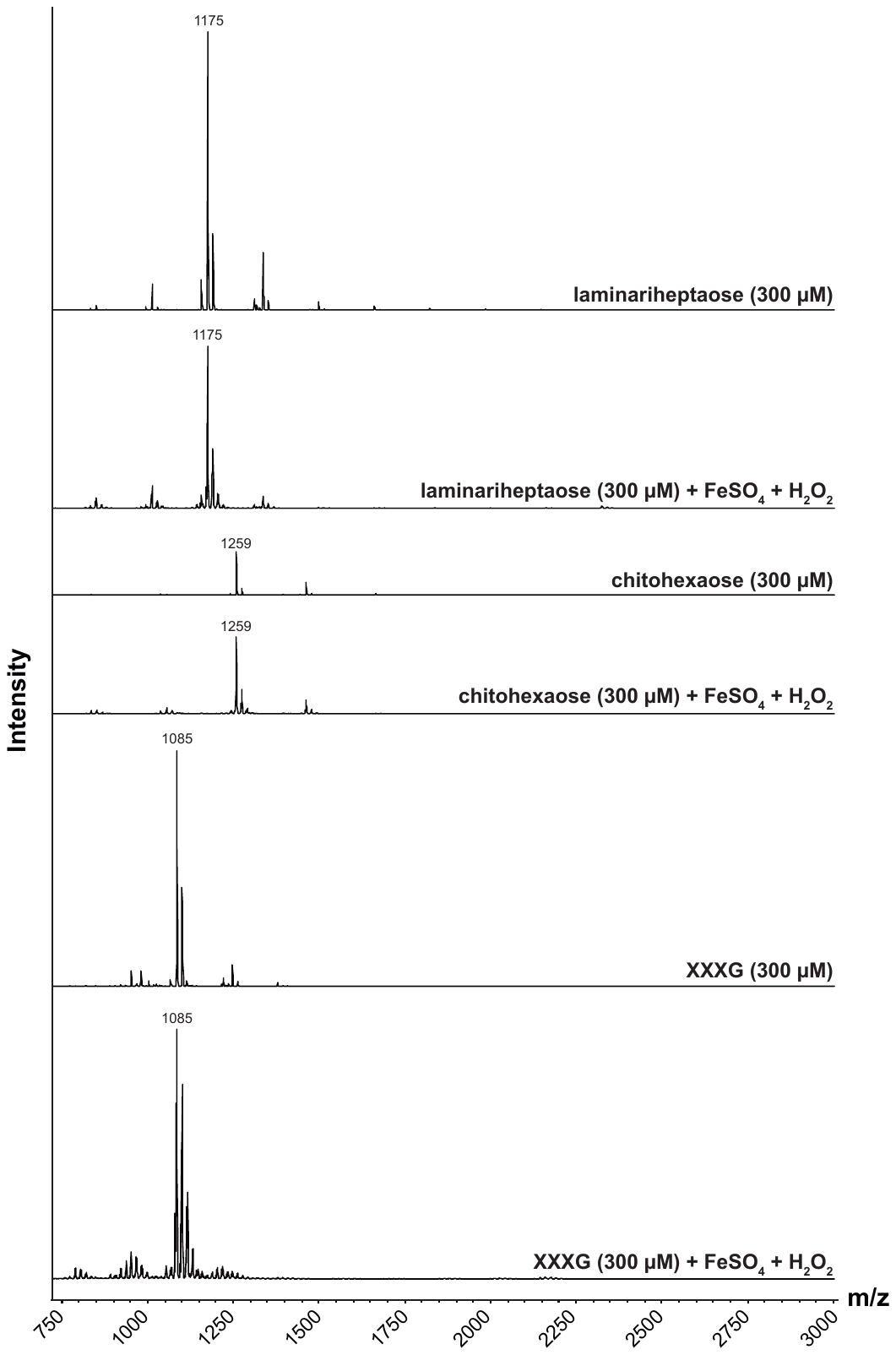

**Figure S12. Glycan controls are not degraded by hydrogen peroxide during Fenton reaction.** Laminariheptaose, chitohexaose and xyloglucan heptasaccharide (XXXG) were tested for oxidative degradation by overnight Fenton reaction (1 mM H_2_O_2_, 100 µM FeSO_4_) followed by MALDI-TOF analysis.

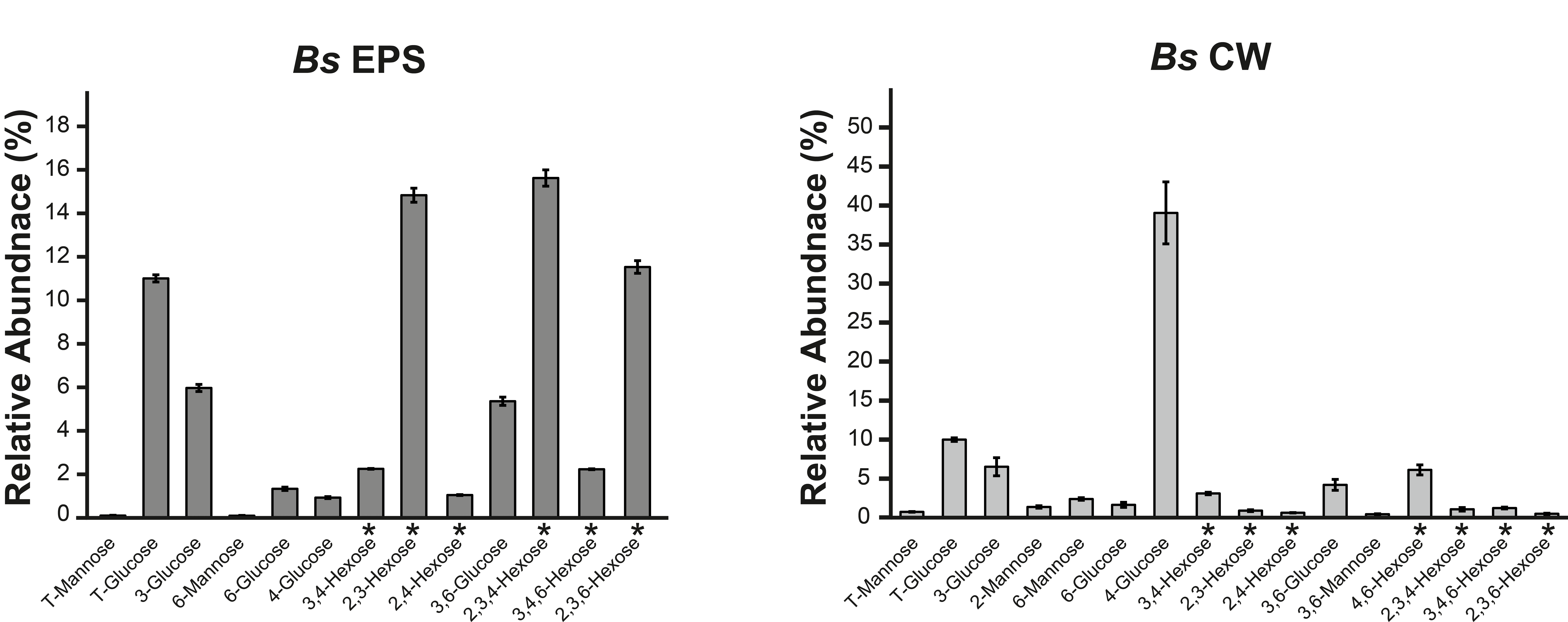

**Figure S13. Glycosyl linkage analysis of *B. sorokiniana* EPS matrix and CW.** 2 mg of ground EPS matrix or the alcohol insoluble residue (CW fraction) isolated from the cultures of *B. sorokiniana* grown in YPD medium was subjected to glycosyl linkage analysis as previously described (Liu *et al.*, 2015). The sugar residues were derivatized into partially methylated alditol acetates and analyzed using GC-MS. The glycosidic linkages were assigned based on the retention time and the mass spectrum available at CCRC spectral database. The annotated glycosyl residues detected in the EPS matrix or CW are expressed as a percentage of total glycosyl peak areas. The other minor unannotated glycosyl residues can be found in the Supplemental Table S6 and S7. Average and standard deviation of three independent biological replicates are presented. *Bs*: *Bipolaris sorokiniana*; CW: cell wall; EPS: extracellular polysaccharide.

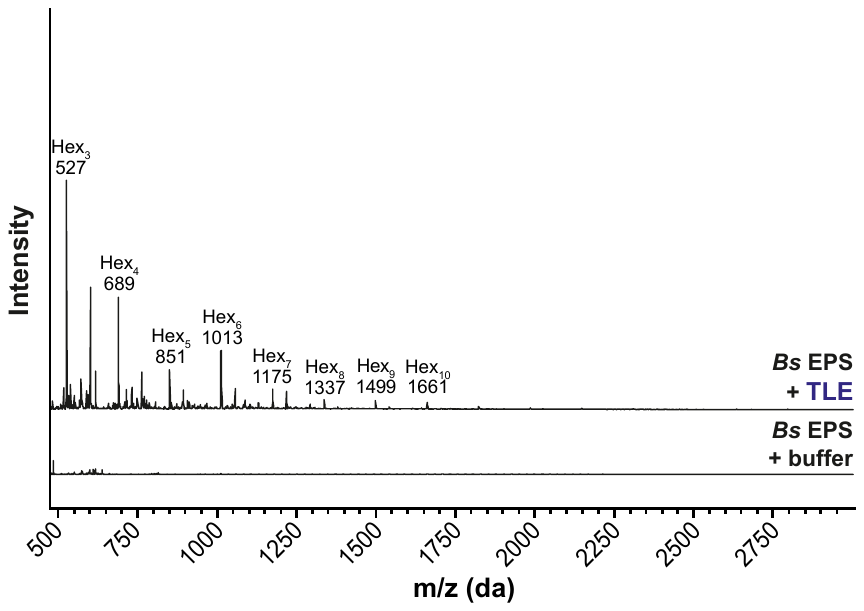

**Figure S14. Analysis of oligosaccharides released from the EPS of *B. sorokiniana* by the action of *Trichoderma* *harzianum* lysing enzymes.** 1 mg of freeze-dried EPS obtained from *B. sorokiniana* was soaked in 2 mM sodium acetate, pH 5.0 at 70°C overnight. The soaked material was incubated with TLE at 40°C for 16 hours. The supernatant fraction from the digested EPS matrix was analyzed by MALDI-TOF. The m/z (M+Na)^+^ Da of oligosaccharides resulting from the digestion of EPS are indicated with their hexose composition. *Bs:* *Bipolaris sorokiniana*; EPS: extracellular polysaccharide; Hex_n_: oligosaccharides with the indicated hexose composition; TLE: *Trichoderma* *harzianum* lysing enzymes.

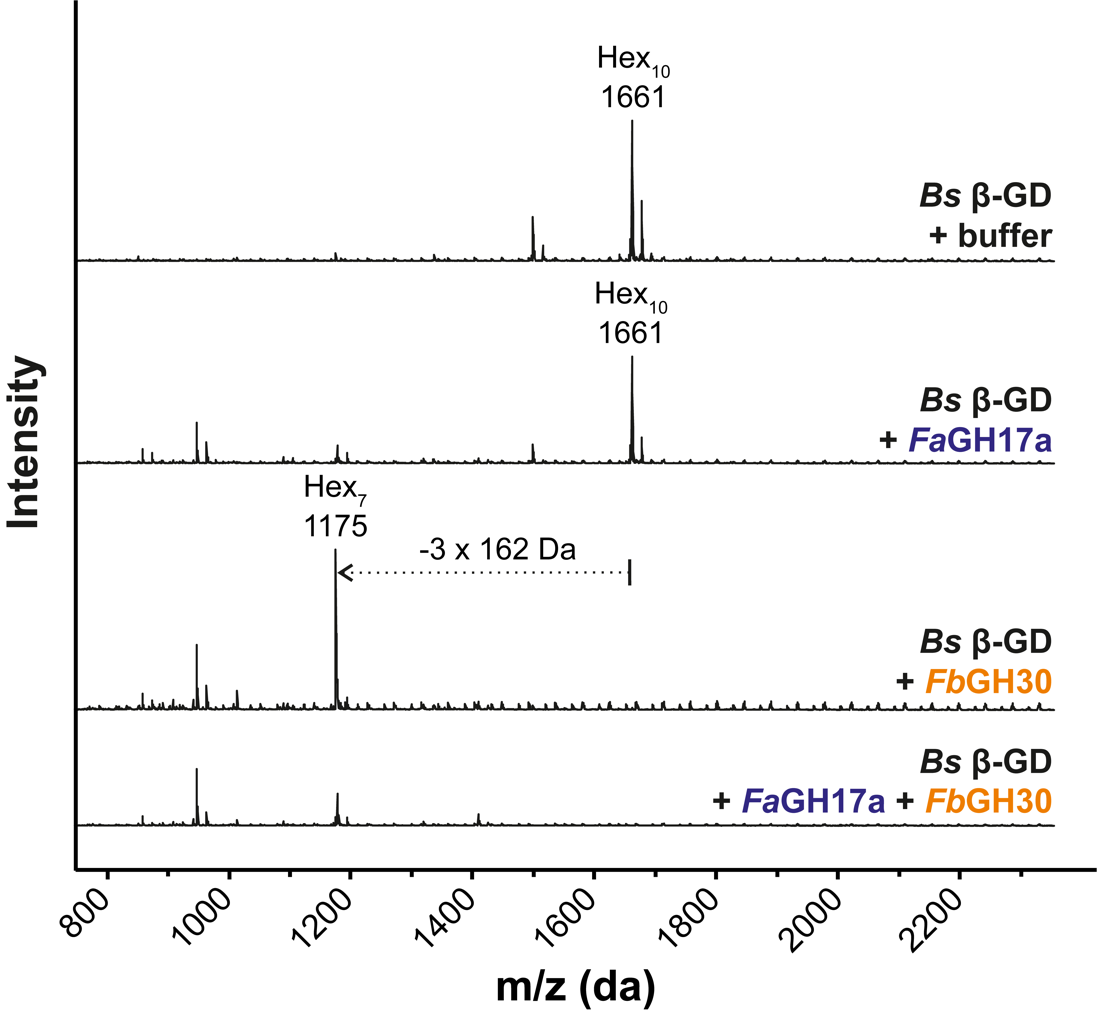

**Figure S15. The β-GD released from the EPS matrix of *B. sorokiniana* consists of a seven unit β-1,3-linked glucan backbone substituted with three β-1,6-glucosyl residues.** β-GD was treated with *Fa*GH17a and/or *Fb*GH30 at 40°C overnight. The digested β-GD fragment was analyzed by MALDI-TOF. The loss of three hexoses (-3x162 Da) as a result of *Fb*GH30 treatment is indicated with a dotted arrow. β-GD: β-1,3;1,6-glucan decasaccharide; *Bs*: *Bipolaris sorokiniana*; EPS: extracellular polysaccharide.

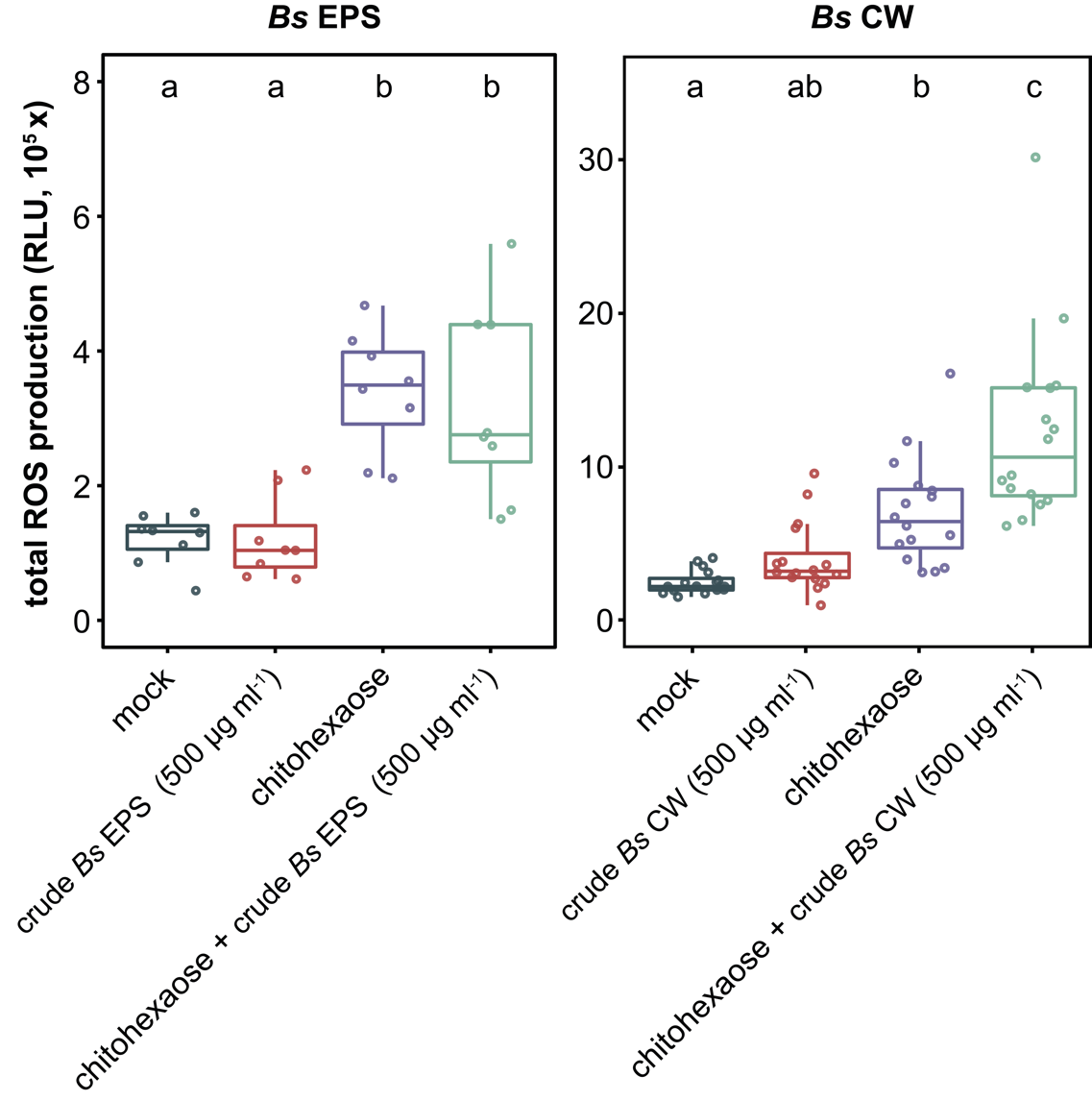

**Figure S16. Mechanically-released fragments from *B. sorokiniana* EPS matrix or CW do not scavenge ROS.** Assays were performed and analyzed as described in Figure S8. *Bs*: *Bipolaris sorokiniana*; CW: cell wall; EPS: extracellular polysaccharide; RLU: relative light units; ROS: reactive oxygen species.

**References:**

**Liu L, Paulitz J, Pauly M**. 2015. The Presence of Fucogalactoxyloglucan and Its Synthesis in Rice Indicates Conserved Functional Importance in Plants1[OPEN]. Plant Physiology **168**, 549-560.

**Wawra S, Fesel P, Widmer H, Neumann U, Lahrmann U, Becker S, Hehemann J-H, Langen G, Zuccaro A**. 2019. FGB1 and WSC3 are in planta-induced β-glucan-binding fungal lectins with different functions. New Phytologist **222**, 1493-1506.
